# Digital PCR quantification of DNA, RNA and extracellular microRNA of mouse oocytes

**DOI:** 10.1101/2021.06.03.446991

**Authors:** Joan Xiaohui Yang, Xin Yuan Zhao, Dexi Bi, Qing Wei, Citra Mattar, Joy Yan Ling Pang, Yie Hou Lee

## Abstract

Despite numerous advances in *in vitro* fertilization (IVF) techniques since its first success in 1978, almost half of the patients treated remain childless. The multifactorial nature of IVF treatment means that success is dependent on variables, including the quality of oocytes. Therefore, new technologies are needed to objectively and quantitatively examine how each oocyte can be selected or optimized to achieve for the best possible outcomes for patients. Here, we report an optimized digital polymerase chain reaction (dPCR) for direct absolute quantification of nucleic acids within 3.5 h without the need for sample extraction or purification. Using individual oocytes, the developed method demonstrated absolute quantification with a linear dynamic range of 0.65 – 33 copies/µL (*r*^*2*^=0.999), high accuracy and excellent reproducibility of <10% relative standard deviation. The method then identified the variable expression of *Gapdh* (0.72-16.95 copies/oocyte), *Hprt1* (1.05-19.05 copies/oocyte) and *ATPase 6*, (5.55-32358.15 copies/oocyte) in ovaries even from the same mouse. Finally, dPCR was used to validate extracellular microRNAs from oocytes incubated with a toxic unsaturated very-long chained ceramide. This study therefore shows the feasibility of dPCR for the rapid and sensitive absolute quantification of DNA/RNA and extracellular miRNA for the study of oocytes.

## INTRODUCTION

The use of assisted reproductive technology (ART) has risen steadily, with more than 8 million babies born from *in vitro* fertilization (IVF) and intracytoplasmic sperm injection (ICSI) since 1978 [1]. With an average of 2.7 treatment cycles required before pregnancy [2] and ART pregnancy rates stabilizing at 36%, improvements are been sought to improve ART outcomes. While ART offers the highest success rate in achieving pregnancy for infertility couples, the take-home baby rate remains disappointingly low [3]. The technology of IVF consists mainly of pick up, selection and insemination of oocytes, embryo culture and their transfer into recipient’s womb. Before any further manipulation of oocytes their quality must be accurately evaluated as it has direct impact on the monospermic fertilization, early development, establishment and maintenance of pregnancy. The main challenge related to selecting high quality oocytes is that developmentally incompetent oocytes may exhibit the same morphologies as the good ones.

The oocyte is a unique and highly specialized cell that supports cellular homeostasis, metabolism, and cell cycle progression in the early embryo, as well as being necessary for creating, activating, and controlling the embryonic genome. During oogenesis, under the instructions of eight transcriptome factors, each oocyte undergoes homologous recombination of paternal and maternal genomes reciprocally exchanging DNA between homologous chromosomes to generate crossovers [4]. Hence, each germ cell has a unique genome that contributes to genetic diversity, resulting in the natural selection of the most competent oocyte to mature and ovulate during each estrous cycle [5,6]. Disruptions in oogenesis through genetic, environmental factors and gynecological disorders can compromise oocyte quality, leading to arrested development and reduced fertility that affect long-term health of the offspring [7]. A viable, high quality oocyte is therefore a prerequisite for successful fertilization and implantation. Oocyte maturation also requires extensive signalling between germ cells and their surrounding theca cells, mural granulosa cells, cumulus cells, all secreting biological factors within the ovarian follicular microenvironment.

Recent studies have revealed the presence of cell-secreted microvesicles and exosomes in follicular fluids, of which extracellular microRNAs (exmiRNAs) are the most abundant constituents [8–10]. miRNAs are a class of small non-coding endogenous RNAs of 19-22 nucleotides that act as post-transcriptional regulators of gene expression [11]. Mature miRNAs are loaded onto the Argonaute (Ago) protein to form the miRNA-induced silencing complex (RISC). miRNA directs RISC to its target mRNA through Watson-Crick base pairing, promoting mRNA destabilization, degradation and translational inhibition [12]. Dysregulated expression of miRNAs such as *miR-206, miR-574, let-7e-5p, miR-29a-3p, miR126-3p, miR-136-5p, miR-192-5p, miR-203a-3p* have frequently been related with oocyte development and aging [13–16]. Our expanded understanding of the molecular determinants of oocyte quality and how these determinants can be disrupted can reveal new insights into the role of oocyte functions in development and improve oocyte development. Therefore, the ability to conduct accurate and sensitive oocyte genetic analysis is critical for gaining insights to oocyte maturation, development, and aging, serving as a basis for understanding infertility disorders and improving fertility outcomes.

In recent years, digital PCR (dPCR) has gained popularity in quantifying genetic material [17–19]. The technique of partitioning allows for high signal-to-noise ratio, high sensitivity to quantify absolute target genes, and being calibration free, are major advantages of dPCR over that of quantitative PCR (qPCR) [20,21]. In addition, dPCR overcomes the shortcomings in qPCR, such as the need for the normalization of the target gene-of-interest to endogenous reference or housekeeping genes, and the need for extraction, both of which result in inaccurate quantification and hampers inter-laboratory comparability of data [22,23] [24]. While direct DNA quantification from plasma or serum using dPCR was previously shown [21,25–27], one step direct detection and RNA quantification in oocytes has not yet been successfully shown. Single oocyte places a physical limitation on RNA isolation due to its minute cellular and genetic volume, as well as the risk of loss. In this study, we optimized a dPCR protocol that is able to provide an absolute quantification of messenger RNA (mRNA) from single oocytes without the need for sample extraction or purification, *i*.*e*. direct quantification. Further, we demonstrated the non-invasive application of dPCR in quantifying extracellular miRNAs in ceramide (Cer) murine oocytes exposed to ceramide (Cer) [28].

## METHODS

### Chemicals

Chemicals, reagents and instruments were from Thermo Scientific unless otherwise stated.

### Murine Oocyte and Lung Harvesting

BALB/c female mice were maintained in a temperature and light controlled, pathogen-free space (24°C, 14 h light/10 h dark cycles) with *ad libitum* food and water. Balb/C female mice (4-8 weeks old) were sacrificed by cervical dislocation and the ovaries were harvested under NUS IACUC protocol R15-1160. The ovaries were collected in oocyte manipulating media [alpha Minimum Essential Medium alpha (αMEM) + GlutaMax media (Gibco, Life Technologies, Carlsbad, CA) supplemented with 25 mM HEPES (Gibco)] and punctured repeatedly with a 30-guage needle to release the cumulus-oocyte complexes (COCs). The COCs were the collected using stripper tips (Origio, CooperSurgical, Charlottesville, VA) with an internal diameter (ID) of 100 μm attached to the STRIPPER® micropipetter (Origio), and place in fresh manipulating media. The surrounding cumulus cells were removed mechanically using ID 75 μm stripper tips. Denuded oocytes were transferred to fresh media and healthy denuded germinal vesicle (GV) oocytes were collected based on their morphology: (i) presence of GV, (ii) oocyte size and shape, and (iii) perivitelline space (PVS). Batches of denuded GV oocytes were rinsed twice in DEPC water + 1% SUPERase in RNase Inhibitor (Ambion™). Several types of buffers were evaluated as part of the dPCR optimization process, namely, RNAlater, DEPC (Invitrogen), DPBS (Lonza), DPBS with RNase inhibitor (SUPERase In RNase Inhibitor), and two types of oocyte culture media: αMEM and *in vitro* maturation (IVM) media [SAGE oocyte maturation media (CooperSurgical, Trumbull, CT)]. 2 μL of buffer was pipetted onto a culture dish and a single denuded GV oocyte was transferred to the droplet. The oocyte and droplet of buffer was then transferred to a PCR tube (total volume of 6 μL) and stored at −80°C until further analysis. As part of the method optimization process, murine lungs were harvested. Lung tissues were washed in PBS and separated into different tubes with the respective buffers. The tissues were then homogenized using Qiagen TissueLyser II at 1 min, 30 Hz, 3 cycles with TissueLyser 5 mm beads. Homogenized tissues were stored at −80°C before analysis (within one week).

### Oocyte Culture Supernatant

Oocyte culture supernatant was collected as previously described [28]. Briefly, groups of 25 denuded GV oocytes were cultured in 30 μL of IVM media supplemented with 25 mIU/mL pregnant mare’s serum gonadotropin (PMSG; Sigma, St. Louis, MO) and 25 mIU/mL human chorionic gonadotropin [hCG (Pregnyl); MSD, Oss, The Netherlands]. The culture drops were layered with mineral oil and matured for 14 h at 37°C in a humidified atmosphere of 5% CO_2_. 60 ng/mL C_24:1_-Cer was spiked into the IVM media of the Cer group. After 14 h, the culture supernatant was collected and stored at −80 °C.

### cDNA Conversion

To convert mouse lung mRNA to complementary DNA (cDNA), a mastermix which comprised of Moloney Murine Leukemia Virus Reverse Transcriptase (M-MLV RT), M-MLV RT 5 × Reaction Buffer, RNAsin Ribonuclease inhibitor, random primers, dNTP mix and RNAse-free water was prepared (**Table S1**). Reverse transcription (RT) was ran at 37°C for 60 min. cDNA concentrations were analyzed using a NanoDrop 2000 spectrophotometer.

### Quantitative Real-Time Polymerase Chain Reaction

Primers for mRNA quantification were designed to bind to non-adenlyated mRNA sequences. Primer sequences are shown in **Table S2**. Each reaction mix consists of 150 nM forward and reverse primers and 1× SYBR Green PCR Master Mix (Applied Biosystems™), 3 μL cDNA sample and PCR grade water topped up to 15 μL (**Table S1**). The reaction mix were then ran at the following cycling conditions: (i) polymerase activation – 95°C for 5 min, (ii) denaturing / annealing & extension – 95°C for 50 s and 58°C for 90 s for 40 cycles. Amplification of the cDNA were analyzed using the Applied Biosystems 7500 Fast Real-Time PCR system. The threshold cycle (CT) was used to determine the relative abundance of the targets.

### Digital Polymerase Chain Reaction

Commercial chips for sample partitioning and matched fluorescence reader for endpoint detection were used (Clarity dPCR system) [26]. For dPCR of mouse lung cDNA, 150 nM of forward and reverse primers and 1× SYBR Green PCR Master Mix, 1× Clarity JN solution, 3 μL DNA sample and PCR grade water topped up to 15 μL. The samples were then delivered onto the Clarity high-density chips using the Clarity auto loader, where it was sub-divided into 10,000 partitions. The partitions were then sealed with the Clarity Sealing Enhancer and 245 μL of Clarity Sealing Fluid was added to each tube. The tube strips were transferred to a thermal cycler for PCR amplification using the following parameters: (i) polymerase activation: 95°C for 5 min, (ii) denaturing / annealing & extension: 95°C for 50 s and 58°C for 90 s for 40 cycles, (iii) final hold: 70°C for 5 min. The ramp rate is kept at 1°C/s throughout the cycling. The tube strips were allowed to cool to room temperature and transferred to the Clarity Reader which detects fluorescent signals from each partition concurrently. Imaged fluorescent signals are shown in **Figure 1A**, wherein positive and negative partitions are displayed as green and blue dots, respectively. In partitions that did not receive any reaction mixture are displayed as black backgrounds. The data were analysed with the Clarity software (version 2.1). Poisson statistics is used to determine the gene copy number per microlitre of sample based on positive signals. Data are expressed as DNA copy number per microlitre of sample or per sample itself. No template controls (NTC) were used as negative controls. From loading of samples to completion of data acquisition, the entire dPCR assay took approximately 3.5 hours.

**Figure 1.**
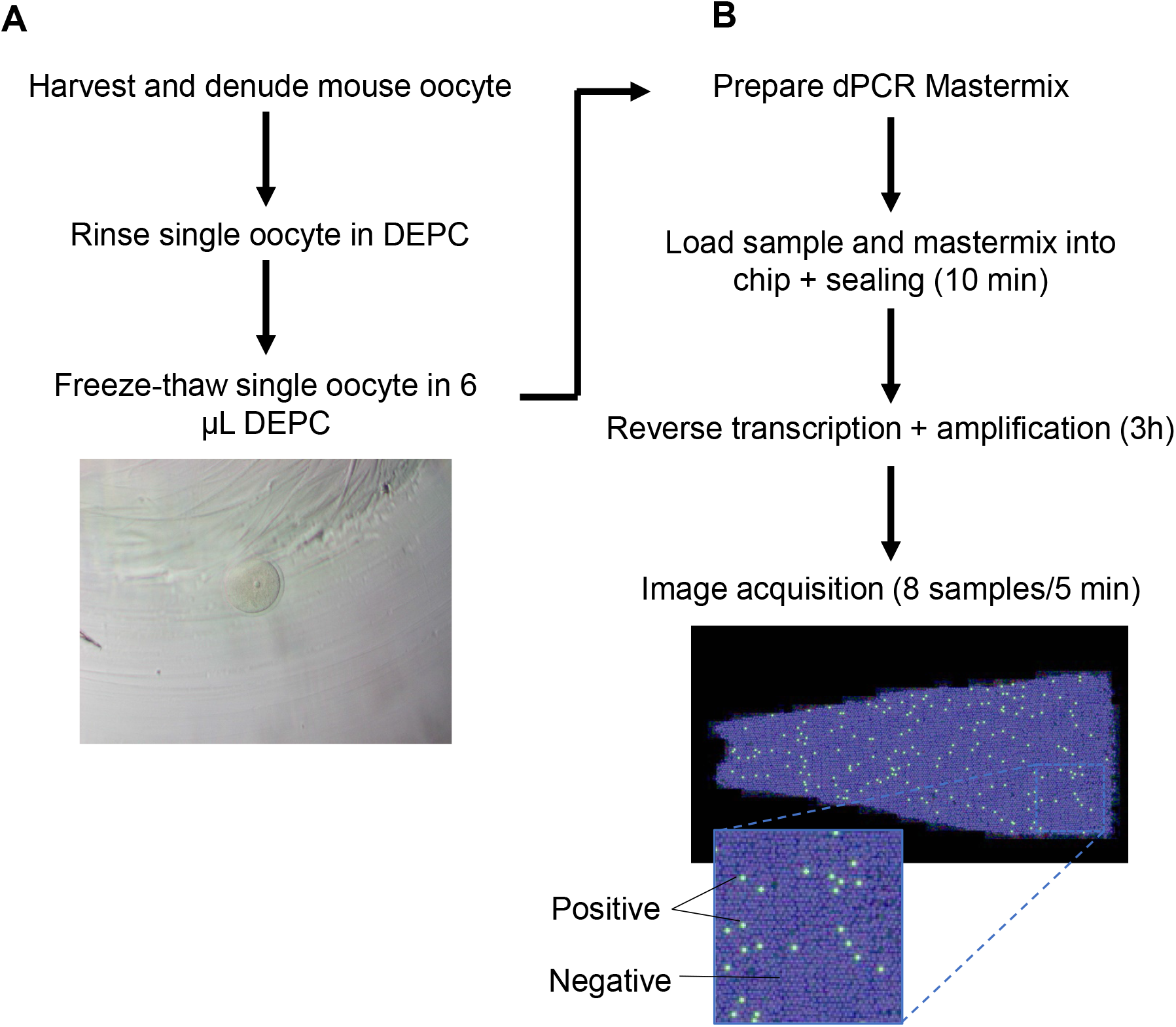
Workflow of direct digital PCR (dPCR) protocol and optimization. (A) Single oocytes were harvested, lysed by freeze-thawing, and proceed to perform (B) one-step RT-dPCR as detailed in text. The entire workflow took from oocyte freeze-thaw to image acquisition was ∼ 4 h.

### Digital PCR - One-Step mRNA Reverse Transcription

This protocol utilizes direct detection of the transcripts, eliminating the need for the purification of mRNA (**Figure 1B**). Assessment of compatible buffers was conducted using *GAPDH* as the model gene given its ubiquitous expression for the evaluation of the buffers. Firstly, 1 ng/µL of total DNA from mouse lung tissue was split and suspended in six different buffers/solutions, namely: RNAlater, DEPC, DPBS, DPBS with RNAse inhibitor and oocyte culture media, alphaMEM and IVM. The most optimal buffer, as evaluated based on readout of *Gapdh*, then went into a second round of buffer compatibility evaluation using mouse oocytes. DEPC was determined as the most compatible buffer. Single oocytes in PCR tubes were thawed and incubated with 0.2 mg/mL of DNase I (Roche) at 37°C for 30 min on a shaker. The reaction mastermix were then added to the oocyte samples directly. Each mastermix consisted of 150 nM forward and reverse primers, 1× QuantiNova RT Mix (Qiagen), 1× QuantiNova SYBR Green RT-PCR Master mix, 1× Clarity™ JN solution and PCR grade water topped up to 15 μL. The reaction mix were loaded onto the chips and sealed before transferring to a thermal cycler. Prior to thermal cycling (as detailed in above for DNA dPCR), a RT cycle of 50°C for 10 min was performed. Data are expressed as mRNA copy number per oocyte.

### Digital PCR - MicroRNA Reverse Transcription

miRCURY LNA miRNA PCR Kit (Qiagen) was used to carry out RT-dPCR on the oocyte supernatant miRNA. To run RT, the supernatant was mixed with 5× RT Reaction Buffer and 10× RT Enzyme Mix and transferred to a thermal cycler (Bio-Rad). Cycling conditions were as follows: (i) RT: 42°C for 60 min, (ii) inactivation: 95°C for 5 min. Following RT in PCR tubes, the miRNA cDNA samples were then added to the dPCR reaction mix), loaded onto the Clarity chips, sealed and amplified as detailed above (in Digital Polymerase Chain Reaction sub-section), with an annealing temperature of 56°C. miRCURY LNA PCR primers (Qiagen) were used for optimal PCR results. Synthetic miRNA sequences (**Table S1**) were designed to optimize the detection of miRNAs in the Clarity platform.

### Profiling of extracellular miRNome

Total RNA was extracted from 20 µL of pooled oocyte culture media using the miRNeasy Serum/Plasma kit (Qiagen) in a final elution volume of 10 µL. Subsequently, the extracted RNA was processed using the miScript Single Cell qPCR kit (Qiagen) according to the manufacturer’s instructions with the following modifications. A total of 5.5 µL extracted RNA was used as starting material for the 3’ ligation reaction. The subsequent 5’ ligation, universal cDNA synthesis, clean-up and universal preamplification steps were performed as per protocol. Sample quality was checked using five control miScript Primer Assays to monitor the 3’ ligation, 5’ ligation, reverse-transcription, preamplification and real-time PCR performance. All samples passed quality control.

The profiling of the 940 most abundantly expressed and best characterized miRNA sequences in the mouse miRNA genome (miRNome) as annotated in miRBase Release 16 (www.mirbase.org) was then performed using miScript Mouse miRNome PCR array (V16.0, 96-well/Rotor-Disc 100; Qiagen) in accordance with the manufacturer’s instructions. Data analysis was performed using the online miScript miRNA PCR Array Data Analysis Tool. Briefly, from 20 μL pooled mouse oocyte culture media from untreated (*N*=3), or C_24:1_-Cer-treated individual mice (*N*=3) [28], miRNA was extracted and converted to cDNA as described above. Next, the cDNA was diluted 1:10 in RNAse/DNAse free water and used as a template for miRNA PCR array amplification. Before that, the PCR array plate was sealed and centrifuged at 1,000× g for 1 min at RT to remove air bubbles that may interfere with the PCR amplification process. Then, PCR reactions were performed using a thermo cycler (Bio-Rad, Hercules, CA, USA). The reaction mixtures were incubated at 95°C for 10 min to activate the HotStart DNA Taq polymerase, followed by 40 cycles of 95°C for 15 s and 60°C for 60 s, monitored by melt curve analysis. C_t_ values >35 were excluded from the analyses.

### Oocyte mRNA-sequencing

Using mRNA from the same oocytes from untreated (*N*=3), or C_24:1_-Cer-treated individual mice (*N*=3) [28] as with the miRNA profiling, library preparation was carried out according to manufacturer’s instructions for QIAseq FX single-cell RNA Kit (Qiagen) using to 7 µL of cell material as starting material. Libraries were quantified using PerfeCTa NGS Quantification Kit for Illumina (Quanta Biosciences) and pooled into a single library for paired end sequencing on one lane of HiSeq4000 using 151 bp read length. Data analysis was then performed using the STAR-RSEM pipeline of GIS’ NGS Pipeline Framework version 2017-10.0

### Bioinformatics Analysis

Prior to bioinformatics analysis, differently expressed mRNAs and miRNAs were identified with the limma package (version 3.10.3) in R (version 3.4.1) [29]. Spearman correlation was performed to identify negatively associated miRNA-mRNA pairs with the miRNA and mRNA expression profiling datasets of all samples as previously described [30] in R and those with *p*<0.01 were subsequently subject to miRNA target analysis with miRWalk2.0 [31], which integrated twelve existing miRNA-target prediction databases. The miRNAs with over ten target genes appearing in ≥6 databases from the miRNA-mRNA pairs were selected and the target genes of each miRNA were subject to Gene Ontology enrichment analysis [32] and KEGG pathway analysis [33]. Gene functions was also cross-referenced with UniProt (https://www.uniprot.org/), when needed. Cytoscape (version 3.7.0) [34] was used to visualise miRNA-mRNA networks.

### Statistical Analysis

Fold change was calculated as the average ratio of normalized molecular tag counts (or relative miRNA expression) between the untreated group and C24:1-Cer-treated group. Numbers greater than 1 indicate upregulated or increased miRNA expression, numbers between 0 and 1 indicate downregulated or decreased miRNA expression, and a fold change of 1 indicates no change. *P*-values were calculated using a Student’s t-test (two-tail distribution and equal or unequal variances between the two samples) on the average delta C_t_ values in each Test Group compared to the Control Group.

## RESULTS

### Direct digital DNA and RNA quantification optimization

The direct DNA and RNA dPCR quantification method in single oocyte is shown in **Figure 1**. The development and optimization of the direct dPCR analysis on the chip-based dPCR system consist of three main steps: (i) buffer selection, (ii) determination of quantification accuracy, and (iii) determination of quantification parameters. Assessment of the compatibility of various buffers was first determined using mouse lung tissues, then mouse oocytes. As shown in **Figure S1A**, DEPC gave the highest signal and was 43-folds higher than the next performing buffer (DEPC: 31.3±3.9 versus IVM: 2.2±0.3 copies/µL *Gadph* DNA). The compatibility of top two performing buffers, DEPC and IVM, for direct dPCR quantification of oocytes was then verified. In DEPC, 55.1±8.6 copies/µL of *Gadph* was detected compared to 26.7±7.1 copies/µL in IVM (**Figure S1B**). Sensitivity of the assay was next evaluated. In serially diluted lung tissue cDNA, a linearity at *r*^*2*^=0.999 across a concentration range over at least three orders of magnitude from 0.1 ng/µL to 10 ng/µL (or 0.65 to 33 copies/µL) was achieved (**Table S3A**). The lower limit of detection (LLoD) of dPCR was comparable to qPCR, the ‘gold standard’ of genetic quantification (0.1 ng/µL or 0.65 copies/µL, *r*^*2*^=0.998, as determined by the C_t_ value >30; **Figures 2A, 2B**). Only signals generated in the partitions that separated were above the threshold of background were taken as positive signals (**Figure S2**). Using *ATPase 6*, the direct dPCR LLoD was 0.77 copies (**Figure 2C**), verifying our sensitivity assessment of the assay. Next, the accuracy of dPCR in quantifying absolute gene copies of target genes was assessed by comparing spiked *E. coli GapA* gene of known gene copies (expected) against the dPCR readouts (observed). In a tested range of 112.5 – 1800 copies/µL, the observed versus expected *GapA* gene copies correlated significantly (*r*^*2*^=0.999) with a gradient (*m*) of 0.969 and demonstrated excellent reproducibility of <10% relative standard deviation (**Figure 2D; Table S3B**). To circumvent the need for DNA isolation and instead, perform direct dPCR analysis, we optimized chemical-free lysis methods. Using GAPDH as the reference gene, we observed that 10 quick freeze-thaw cycles of oocytes in liquid N_2_ performed the best compared to 10 slow freeze-thaw cycles and one freeze-thaw lysis (data not shown). Altogether, we optimized the dPCR method for single oocyte analysis, determined DEPC as the buffer of choice, demonstrated accuracy in detecting *bona fide* absolute gene copies using dPCR, and established the sensitivity limits of the dPCR assay.

**Figure 2.**
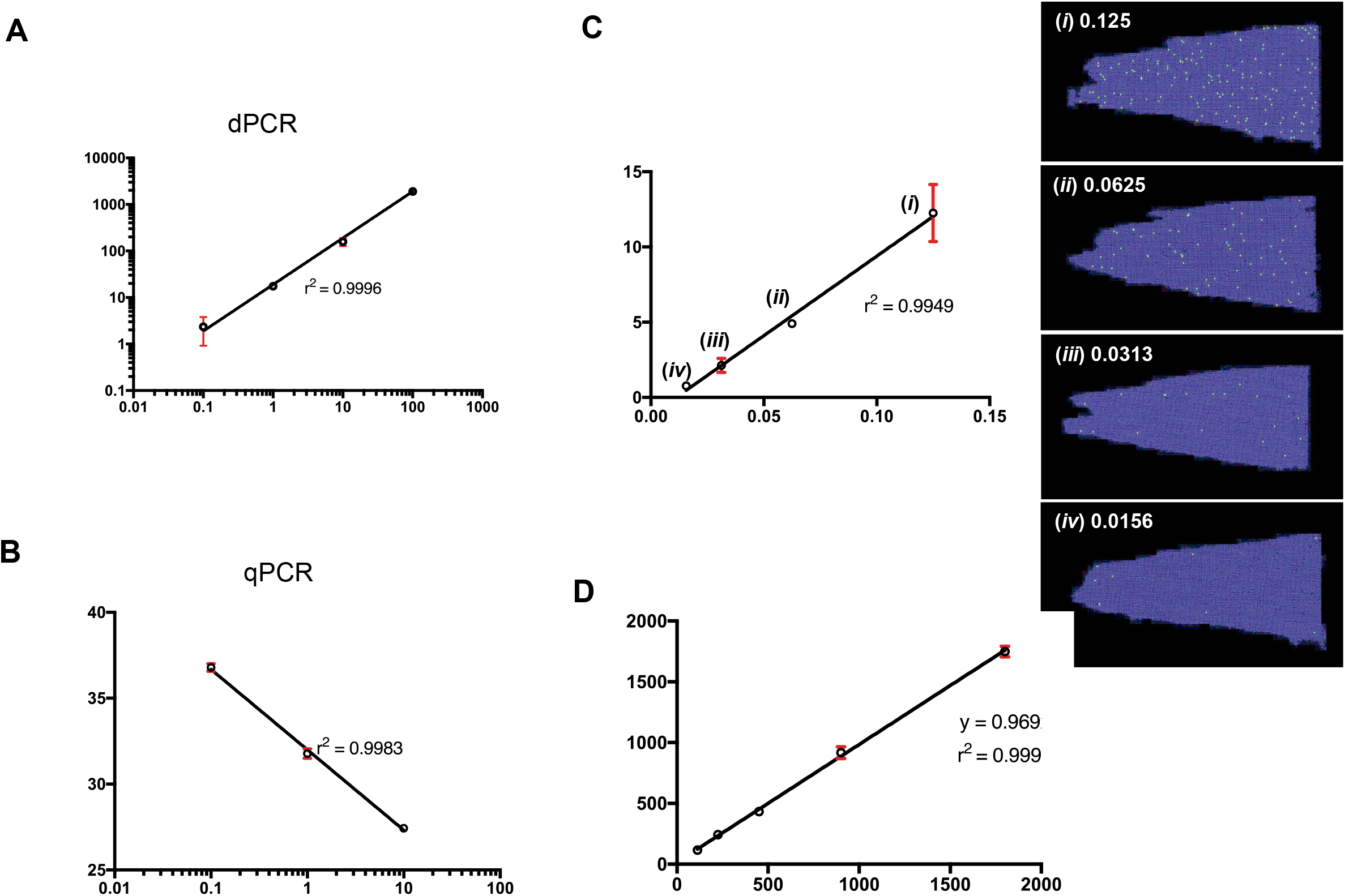
Direct dPCR limit of detection (LLoD). The LLoD of direct dPCR (A) was compared to the LLoD of conventional qPCR (B). Both had LLoD of 0.1 ng/mL. (C) The dilution of a single oocyte was analysed using direct dPCR, showing a linear dilution curve with R^2^ close to 1. By diluting a single oocyte to various ratios and detecting for ATPase6, we were similarly able to show a linear dilution curve down to very low copies of 0.77 copies/uL. The position plots on the right are the representative plots of each dilution. With each dilution, there is a corresponding decrease in the number of positive signals. It is important to note here that the number of dilution of a single oocyte would be dependent on how abundant the gene is expressed in the oocyte itself. (D) Quantification of absolute gene copies of target gene by direct dPCR was assessed by comparing spiked *E. coli GapA* gene of known gene copies (expected) against the dPCR readouts (observed). In a tested range of 112.5 – 1800 copies/µL, the observed versus expected *GapA* gene copies correlated significantly (*r*^*2*^=0.999) with a gradient (*m*) of 0.969.

### Direct digital mRNA analysis of individual oocytes reveals individualized genetic makeup

Having optimized the direct dPCR assay, we tested the assay to perform absolute quantification in single GV oocytes. From the ovaries of five mice, DNA quantification by direct dPCR showed an average of 1067.85 copies/oocyte of *ATPase 6*, 1.20 copies/oocyte of *BubR1*, 5.10 copies/oocyte of *Emil1*, 27.90 copies/oocyte of *Pttg1*, 88.20 copies/oocyte of *Zp3* and 131.40 copies/oocyte of *Gapdh* (**Figure 3A**). We verified oocyte lysis by microscopy. Additionally, we rationalized that if an intact oocyte was analysed there would be no amplification. Transcript levels of *Gapdh* and *Hprt1*, commonly used housekeeping genes [22], were measured from GV oocytes obtained from the ovaries of five mice. Measured mRNA copies varied widely, even within ovaries from the same mouse (*Gapdh*: 0.72-16.95 copies/oocyte, range: 16.23 copies/oocyte; *Hprt1*: 1.05-19.05 copies/oocyte, range: 18 copies/oocyte; **Figure 3B**). *ATPase 6*, which has been shown to be highly induced in metaphase II oocytes compared to GV oocytes [35,36], also varied widely from GV oocyte to GV oocyte (5.55-32358.15 copies/oocyte, range: 32352.6 copies/oocyte; **Figure 3C**). Therefore, absolute quantification of these expressed genes revealed individual oocyte natural variation.

**Figure 3.**
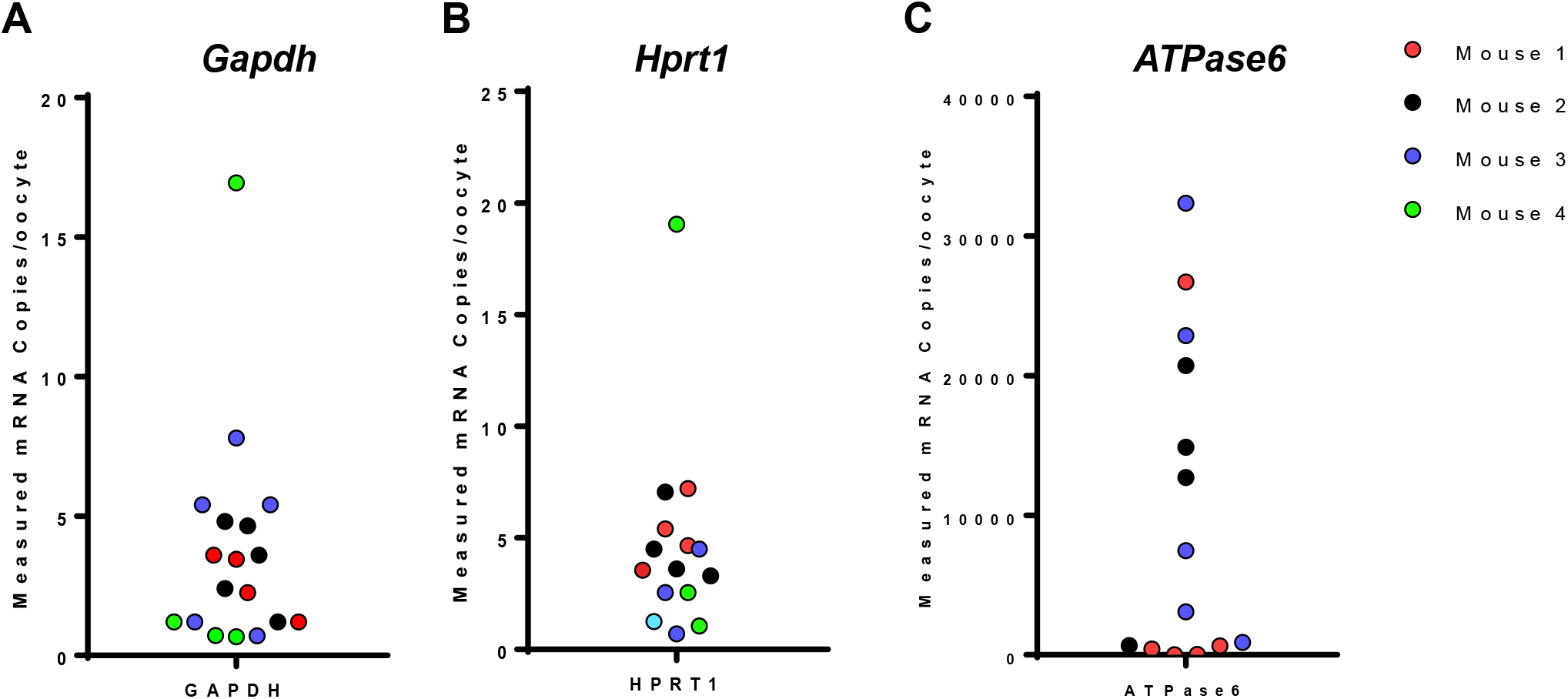
Natural variation of mRNA from individual GV oocytes. Direct dPCR of *Gapdh, Hprt1* and *ATPase 6* mRNA revealed wide variation within ovaries from the same mouse (*Gapdh*: 0.72-16.95 copies/oocyte, range: 16.23 copies/oocyte; *Hprt1*: 1.05-19.05 copies/oocyte, range: 18 copies/oocyte; *ATPase 6*: 5.55-32358.15 copies/oocyte, range: 32352.6 copies/oocyte). Dots of the same colour represent the oocytes obtained from the same mouse. *N*=4 mice.

### Diverse microRNA-mRNA interactions coordinate C_24:1_-Cer oocyte induced maturation inhibition

We extended the dPCR method to probing the extracellular miRNAs (exmiRNome) secreted by oocytes. C_24:1_-Cer increases mitochondrial reactive oxygen species which leads to obtund oocyte maturation rates [28]. Poor maturation of oocytes from MI to MII may be associated with suboptimal interactions of the exmiRNA from poorly matured oocytes with themselves, their microenvironment and surrounding cells [37]. Among 940 miRNAs that were profiled, a total of 7 exmiRNAs were upregulated and 198 down-regulated exmiRNAs in C_24:1_-Cer treated murine oocytes compared to untreated oocytes (log_2_ fold change >2 or <-2, *p*-value <0.05) (**Figure 4A; Table S4**). Among these, three exmiRNAs *miR-380-5p, mmu-miR-1969*, and *mmu-miR-509-5p* were selected for validation by dPCR. Significant reductions in levels of two exmiRNAs, *mmu-miR-1969 and mmu-miR-509-5p*, were validated by dPCR and their copy numbers quantified in the culture media from an independent set of oocytes (**Figure 4B**).

**Figure 4.**
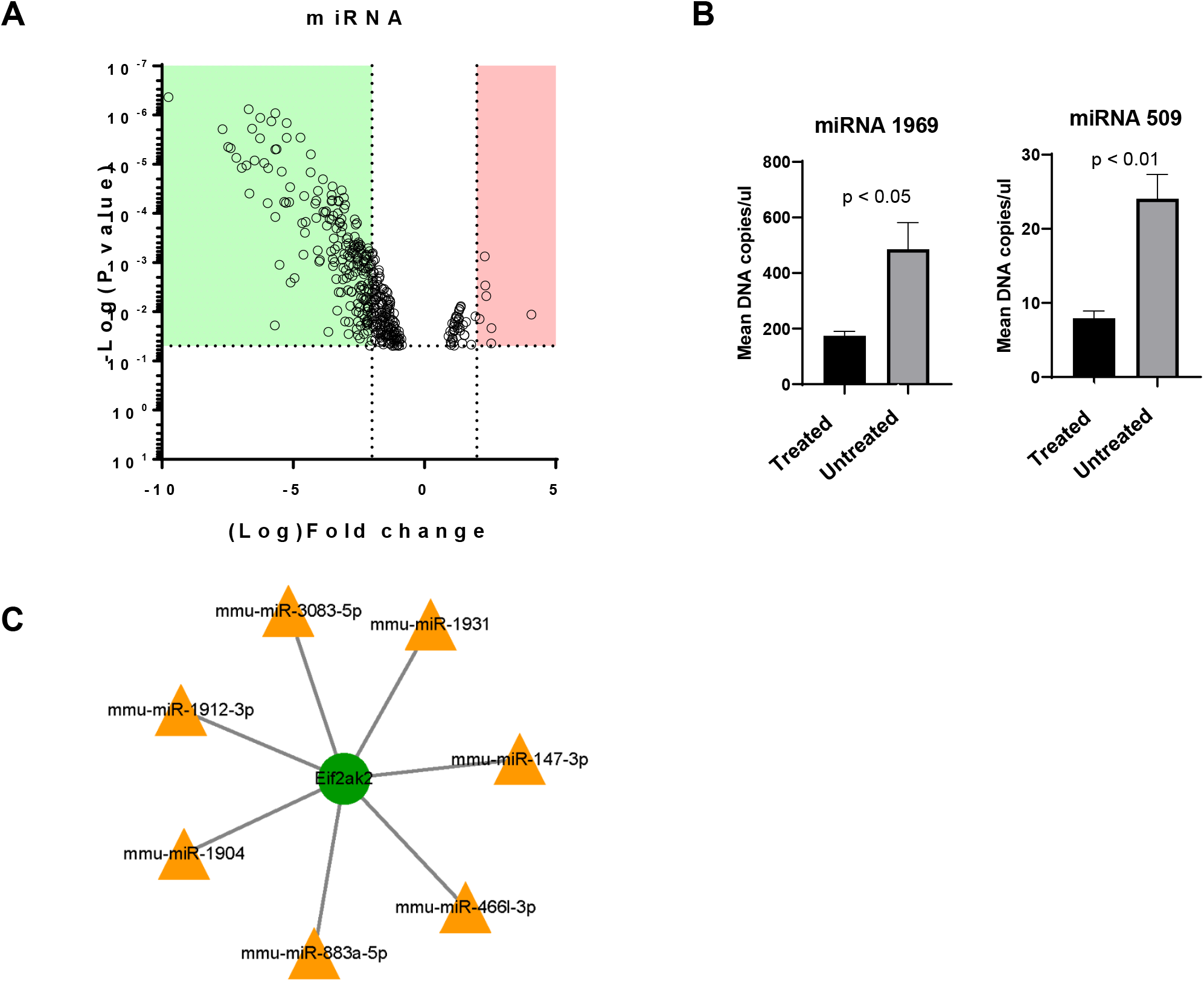
Extracellular microRNA (exmiRNA) profiling and network biology analysis of oocytes upon C_24:1_-Cer exposure. (A) Volcano plot of significantly differential exmiRNA from conditioned media of oocytes incubated with C_24:1_-Cer and vehicle (control) as quantified using miRNome PCR array. 212 miRNAs were upregulated (red) and 151 miRNAs were downregulated (green) in the C_24:1_-Cer treated group. (B) Decrease in miRNA 1969 and 509 were validated by dPCR from an independent set of oocytes (*N*=3). (C) Bioinformatic analysis of miRNA-mRNA pairs of *Eif2ak2*, a gene that play key roles in regulating signal transduction of apoptosis, cell proliferation and differentiation.

To further understand the potential roles of exmiRNA with C_24:1_-Cer exposure, we investigated oocyte miRNA-mRNA target associations. Oocyte transcriptomes were sequenced, and paired miRNA-mRNA profiles were integrated. Two of the validated exmiRNAs, *mmu-miR-1969* and *mmu-miR-509-5p*, were predicted to bind to Bcl2 modifying factor, *bmf*, a member of the BCL2 family that act as apoptosis regulators. In addition, *mmu-miR-509-5p* was predicted to bind to and modulate *Slc22a27*, a sodium-independent organic anion transporter which exhibits high specificity for L-carnitine, a metabolite that improves oocyte maturation when supplemented in culture [38]. Six miRNAs *(mmu-miR-3083-5p, mmu-miR-466l-3p, mmu-miR-883a-5p, mmu-miR-15b-5p, mmu-miR-188-3p* and *mmu-miR-326-5p*) with over ten negatively associated target genes (**Table 1**), consistent with the expected degradation of target mRNAs and/or inhibiting their translation. The target genes participated in different GO biological processes (BPs) or pathways, suggesting that these miRNAs have versatile regulatory roles in oocyte maturation by targeting diverse genes. KEGG enrichment analysis of miRNA-mRNA association revealed glycosphingolipid biosynthesis, fatty acid metabolism, amino acid metabolism and glycan metabolism (**Table 1**). Interestingly, six viral disease-associated pathways were enriched in mmu-miR-466l-3p. Upon further analysis, all pathways had a shared gene, *Eif2ak2*, that play key roles in regulating signal transduction of apoptosis, cell proliferation and differentiation (**Figure 4C**) [39]. Therefore, the results suggest *Eif2ak2*-associated proliferation and apoptotic processes rather than a viral response in C_24:1_-Cer obtunded oocytes. Collectively, diverse miRNA-mRNA interactions coordinate the effects C_24:1_-Cer on obtunding oocyte maturation.

**Table 1.**
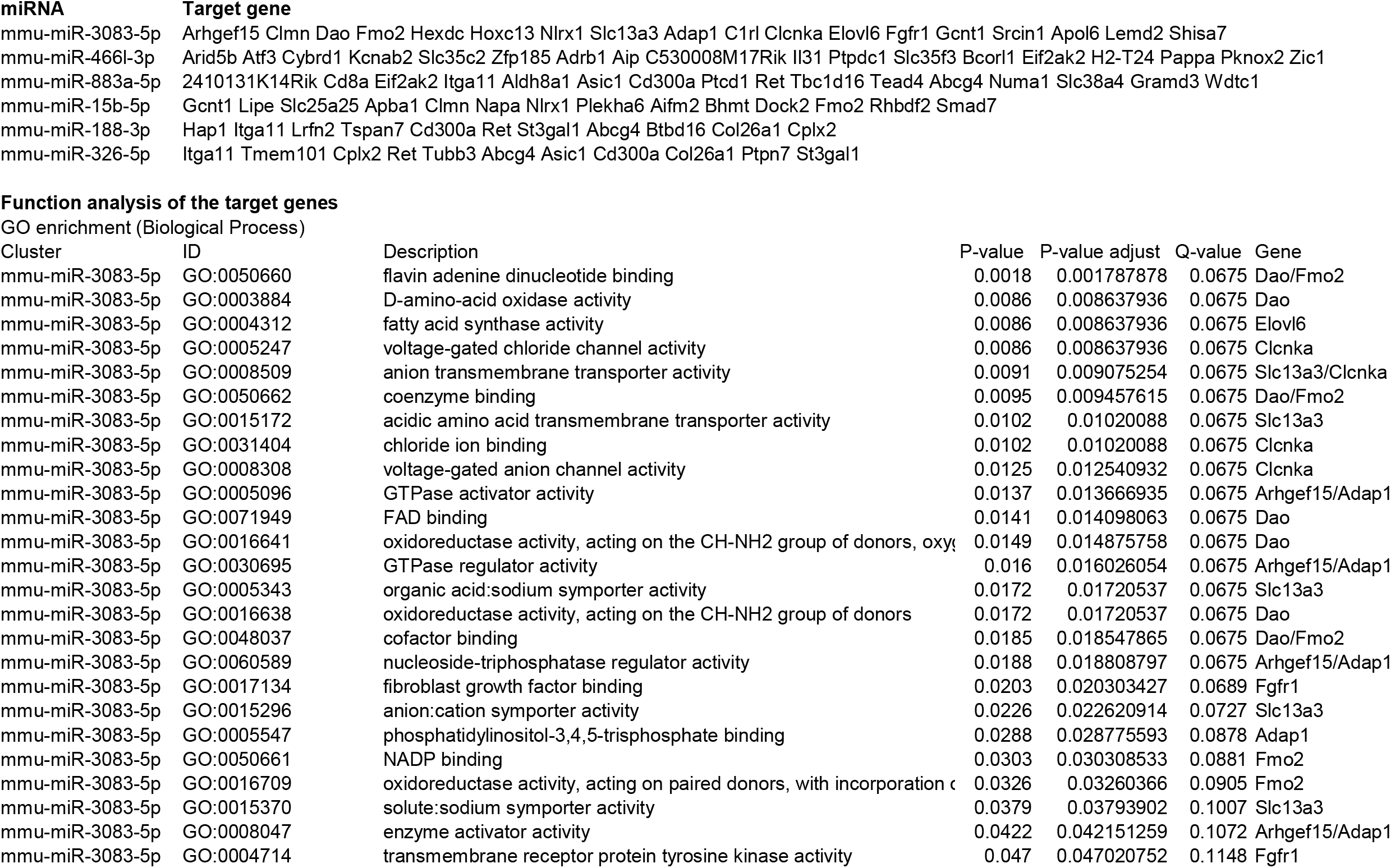
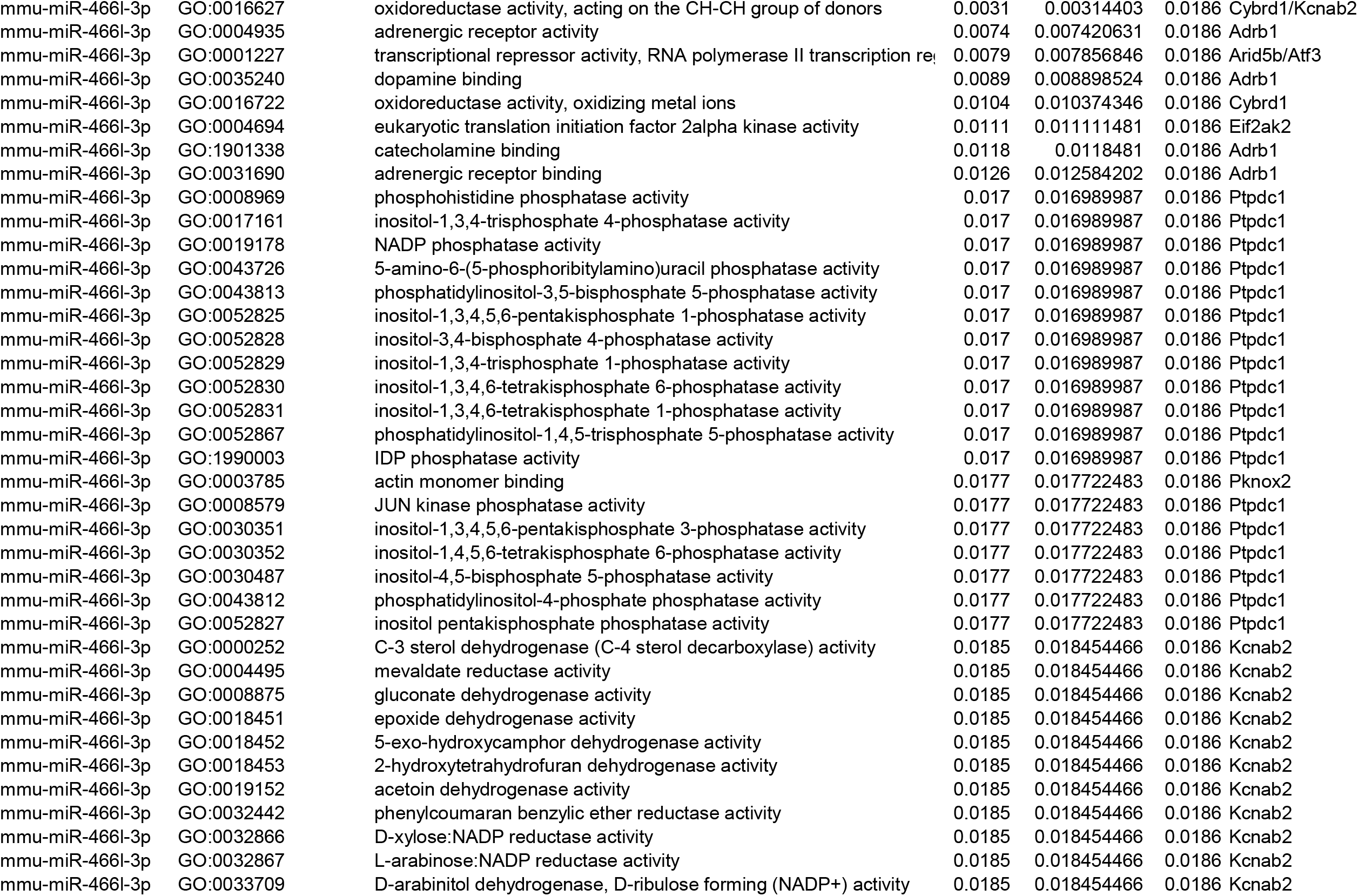
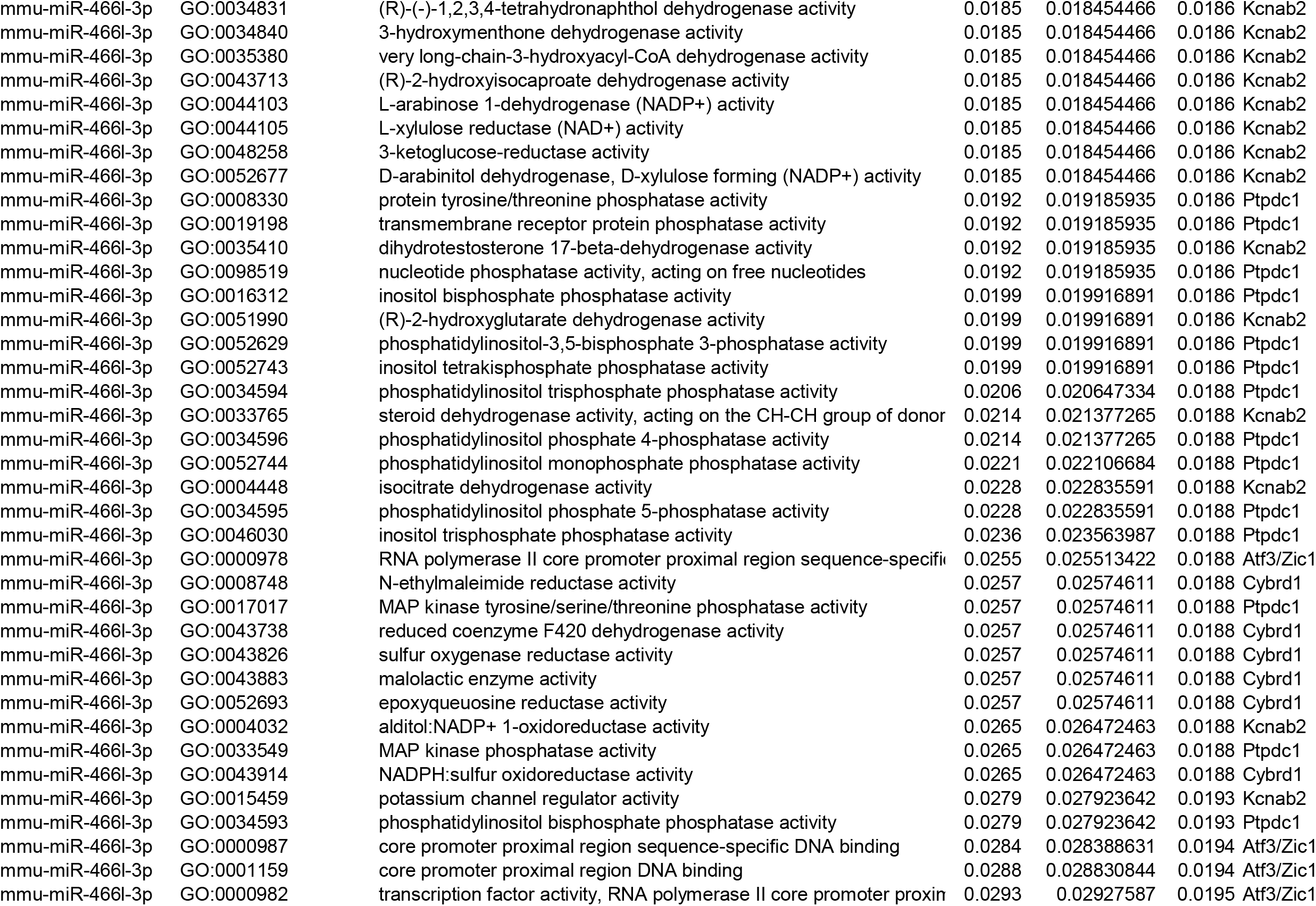
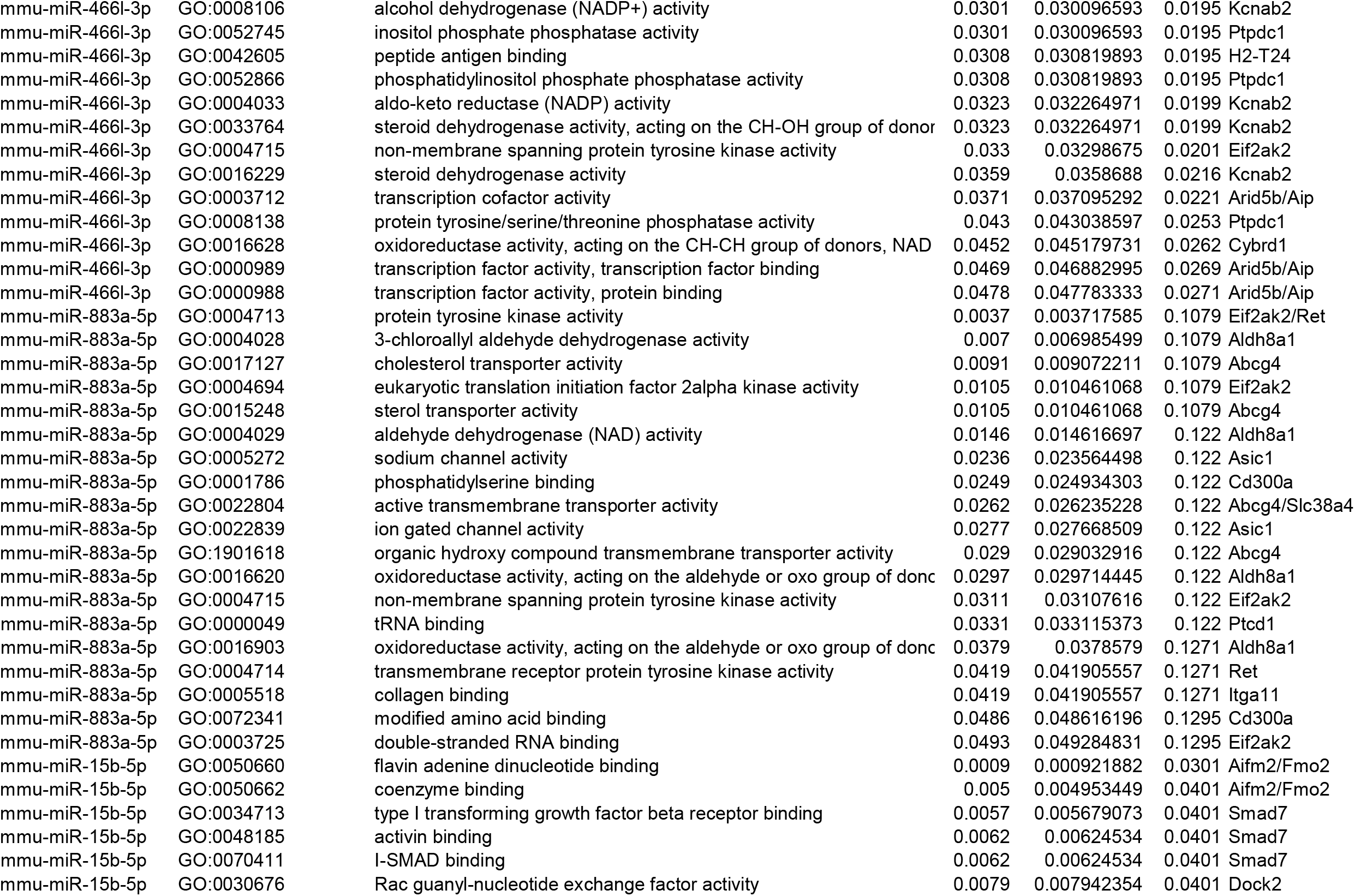
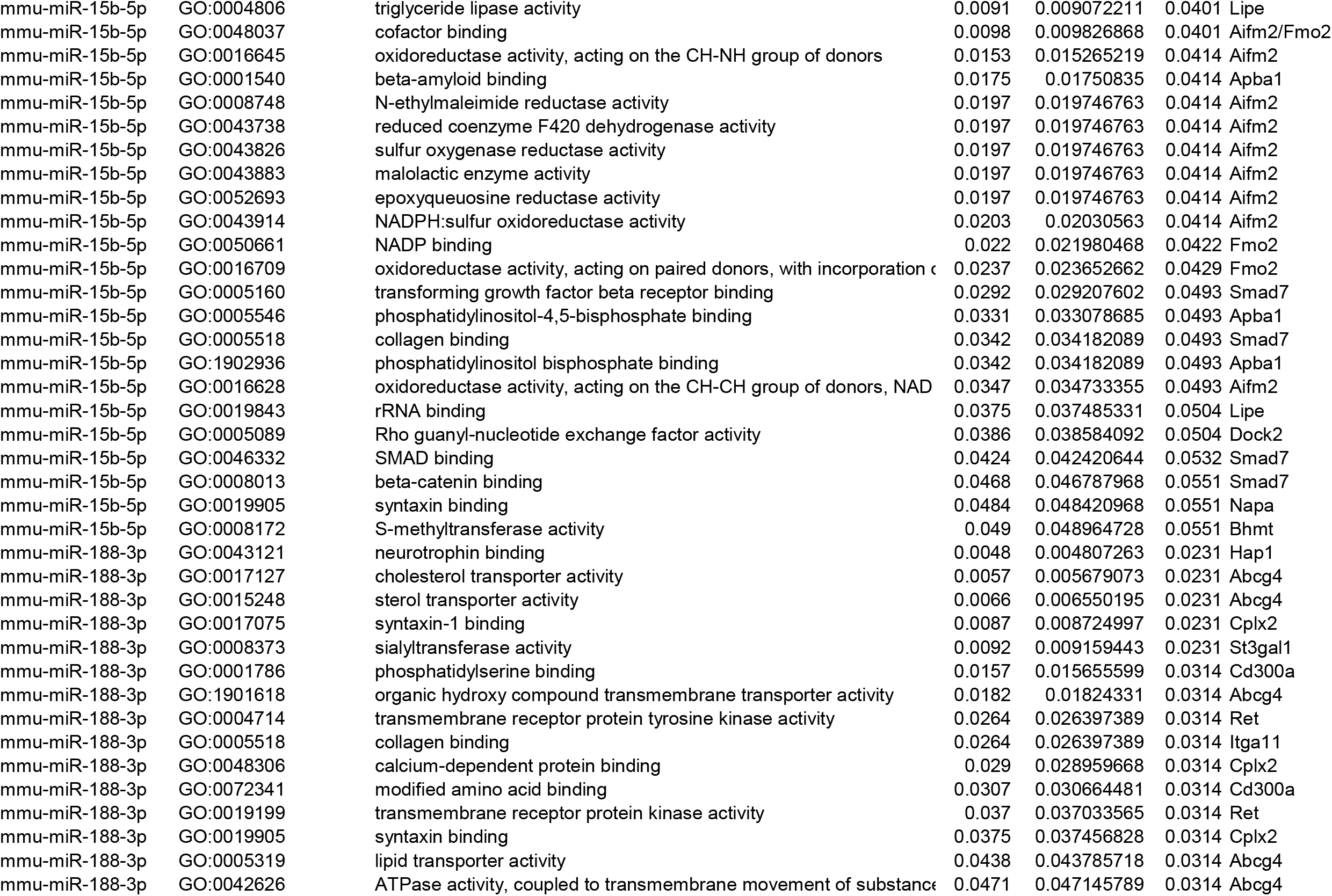
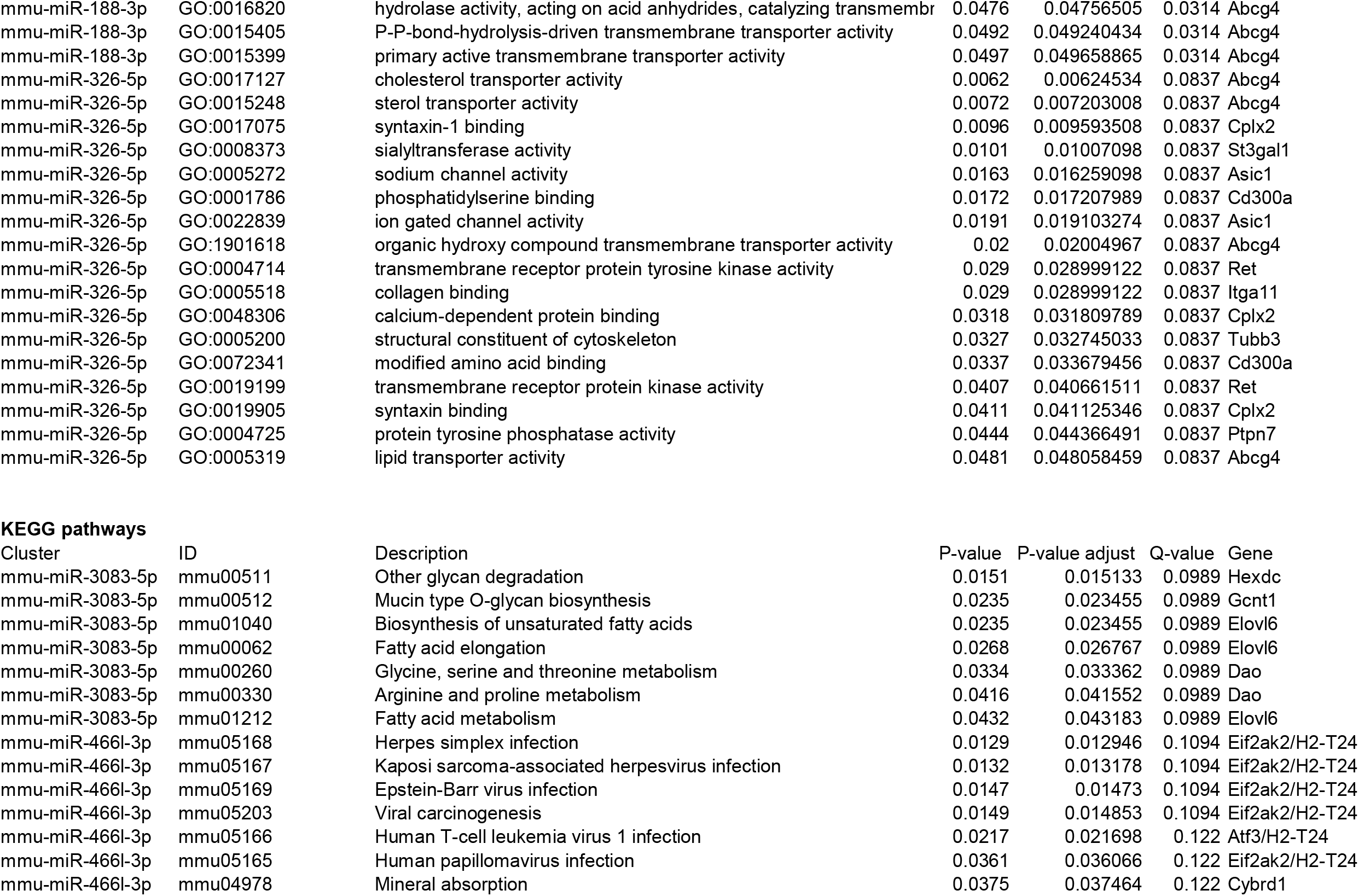
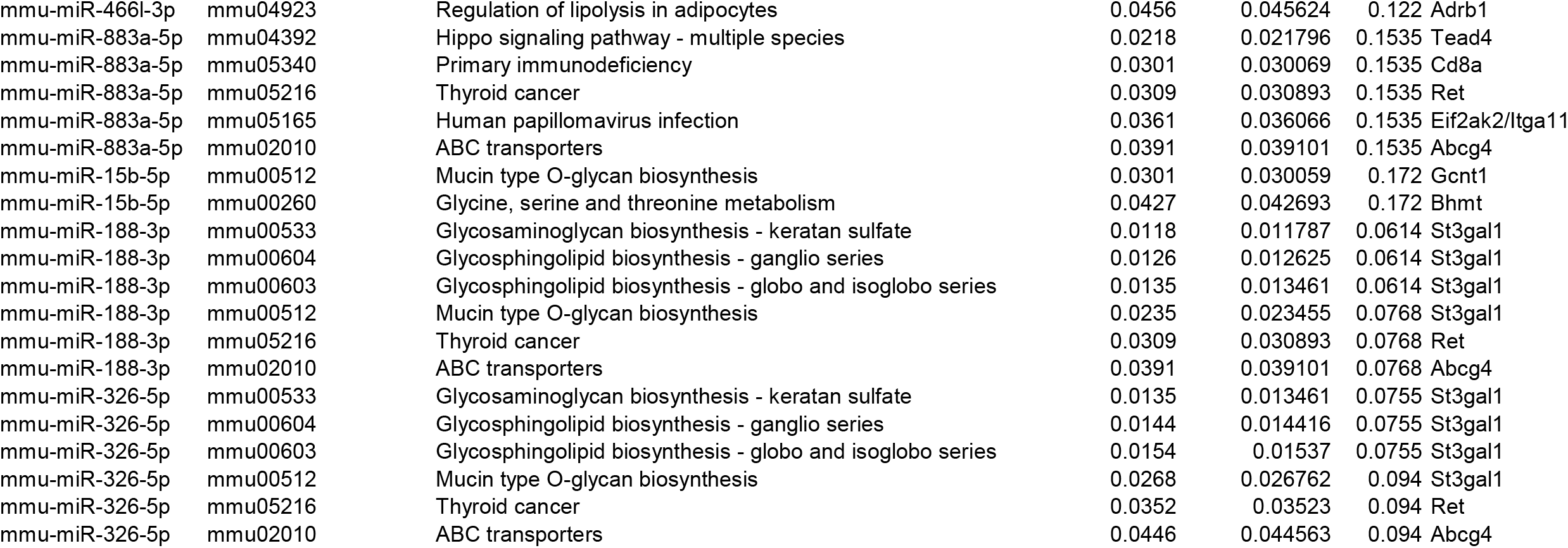
Gene Ontology and KEGG analysis of oocyte miRNA-mRNA associations with C24:1-ceramide.

## DISCUSSION

A high-quality oocyte is a critical prerequisite for successful fertilization and implantation, profoundly suggesting that fertility outcomes might be based on the oocyte variation. Single cell genetic analysis is increasingly investigated across different platforms because of the critical information divulged at the individual cell level [40–43]. In this study, we demonstrate direct dPCR analysis of DNA and RNA from single oocytes without the need for an isolation step, removing the need for extraction or purification of genetic material employed by most techniques [22,40,44–46]. The direct dPCR method was then applied to investigate six genes related to oocyte maturation. In addition to revealing heterogeneous expression of these genes in individual oocytes, this approach provided miRNA-target patterns of how C_24:1_-Cer, a very long chained ceramide previously implicated in endometriosis-associated infertility [28], influences the oocyte transcriptome and prevent oocyte maturation.

In conventional qPCR, housekeeping genes are used for normalization purposes of the targeted gene across different samples, treatment groups or conditions [47–49]. However, the utility of these genes for normalization has been questioned given possible different gene copies varying across samples and conditions [22,50–52]. One major advantage of dPCR is its absolute and precise quantification. In our study, using dPCR we also showed strong correlation between what was measured and what was in the sample. Standard qPCR protocols involve the use of varying amounts of inhibitory substances, consequently leading to quantification inaccuracies and hampering inter-laboratory comparability of data [24]. In contrast, dPCR is more resistant to inhibitory effects of different matrices and is not dependent on calibration and its commutability [53]. Single sperm analysis using a different dPCR platform has been previously reported [54]. In addition, direct dPCR analysis of unpurified plasma or serum has been reported [21]. Major assay advantages in the Clarity™ system used in this study compared with Fluidigm system used in the sperm analysis [54] include reduction in time to analyze samples (<4 h versus >250 h for analysis of 96 reactions), therefore facilitating *bona fide* high-throughput, and reduction in cost. Our assay also allows for straight-forward porting and adaptation of existing qRT-PCR protocols from already established laboratory-customized protocols to systems such as the Clarity™ dPCR platform used herein, as well as QuantStudio 3D Digital PCR System, QIAcuity Digital PCR System, and Droplet Digital PCR System. This is critical as laboratories are likely to have optimized their specific protocols, and easy portability will be favoured. In combination with the absolute and accurate quantification offered by dPCR, rapid and direct detection and quantification of genomic material without minimal loss of sample material and broad applicability is therefore an attractive method for evaluating oocyte quality.

An important observation of this study is the varying amounts of six genes involved in oocyte maturation in individual oocytes. While the maternal transcriptome is reported to be inactive during oogenesis and maturation and reactivate after fertilization, our results are consistent with other studies that certain loci in mouse and Drosophila are transcriptome active [55–57]. A high-quality oocyte is a critical prerequisite for successful fertilization and implantation, profoundly suggesting that fertility outcomes might be associated with oocyte variation. Furthermore, it has been shown that the phenotype of offspring are considerably defined by the quality of the oocytes from which they are derived [55–57]. The variability of genetic material copies of oocytes from the ovaries of the same mouse as shown in this study exemplifies the need to assess oocytes individually. The ability to conduct accurate and sensitive oocyte genetic analysis is critical for gaining insights to oocyte maturation, development, aging, and oocyte production in vitro, serving as a basis for understanding infertility disorders, determining fertility outcomes [58]. In addition, our findings underpin the importance of oocyte selection for assisted reproductive techniques or ART, whereby multiple oocytes are harvested from which one oocyte would be selected for artificial insemination, paving the way for increasing the chance for implantation [59].

Extracellular miRNAs are emerging as signalling molecules and raises the prospect of oocyte-to-oocyte or oocyte-to-neighbouring cell communications such as thecal cells for the development of oocytes. Many target mRNAs carry multiple miRNA sites that overlap or are in close vicinity. Binding of a miRNA to a target site may be either enhanced or repressed by the changing miRNA milieu [60]. Prominent miRNAs that were significantly altered with exogeneous C_24:1_-Cer affected in oocyte maturation (Table S4) is consistent with other studies. *miR-15b* is known to target genes involved in cell cycle regulation and cell death [61,62]. The upregulation of *miR-15b-5p* is implicated in the development of ovarian toxicity [63]. Furthermore, *miR-15b-5p* has been shown to target estrogen-related receptor gamma (*Esrrg*). Decreased expression of *miR-15b* has been associated with the production of mitochondrial-associated ROS [64], congruent with the increase of mitochondrial ROS induced by C_24:1_-Cer [28]. Freudzon *et al*. reported that *mmu-miR-466l-3p* targets GPR3 gene which maintains meiotic arrest in oocytes [65]. The downregulation of *Gpr3* results in premature ovarian aging [66]. In human follicular fluids of mature oocytes, *miR-380-5p* is one of the more abundant miRNAs [16]. Validated decrease in *miR-380-5p* in our study plausible reflects oocyte maturation inhibition and suggest that C_24:1_-Cer leads to *miR-380-5p* downregulation and hence oocyte maturation inhibition. Therefore, *miR-380-5p* offers the possibility of non-invasively selecting the most competent oocyte based on selected miRNA quantification, although further validation will be required.

Future work includes broad mRNA profiling of oocytes to further evaluate degree of oocyte transcriptome heterogeneity and the impact of oocyte transcriptome upon C_24:1_-Cer exposure. Quality and integrity attributes of oocytes and reliable readouts of single oocytes are critical requisites for quantitative analysis of oocyte quality if such a method would to be employed clinically, which will inadvertently impact the success of ART. The identification and validation of exmiRNAs correlated to oocyte maturation in mouse and humans, including *miR-380-5p*, will add confidence to their use as non-invasive biomarkers of oocyte quality. Digital CRISPR, which was also developed on the Clarity dPCR platform [67] can be evaluated and considered for improved detection of exmiRNA in oocytes.

In conclusion, our study optimized a dPCR protocol that can be performed in 3 h, and demonstrated the feasibility to quantify mRNA from single oocytes without the need for sample extraction or purification. Further, we demonstrated the non-invasive application of dPCR in quantifying exmiRNAs oocyte culture media [28], paving the possibility of investigating extracellular miRNAs from blastocysts and embryos [68] but at the single cell level.

## ACKNOWLEDGEMENTS

This work was supported by SingHealth Foundation (SHF/FG560P/2014) and National Medical Research Council, Singapore (NMRC/BNIG/2033/2015).

## Notes

**Disclosure summary:** All authors have nothing to disclose.

### Competing Interest Statement

The authors have declared no competing interest.

