## Supplemental Figures and Tables for "Digital PCR quantification of DNA, RNA and extracellular microRNA of mouse oocytes"

### SUPPLEMENTARY FIGURE LEGENDS

**Figure S1. Optimization of direct dPCR. The dPCR protocol was optimized to test for the most suitable buffer in (A) mouse lung tissues and (B) mouse oocytes.**

**Figure S2. Thresholding above noise (blue dots) determines the positive signals (green dots).**

**Table S1. PCR components**

**Table S2. Primer sequences**

**Table S3. Summary of (A) PCR chacteristics of GAPDH and (B) comparison of expected and observed E. coli GapA concentration as measured using dPCR.**

**Table S4. Extracellular microRNA by GV oocytes incubated with C24:1 Ceramide**

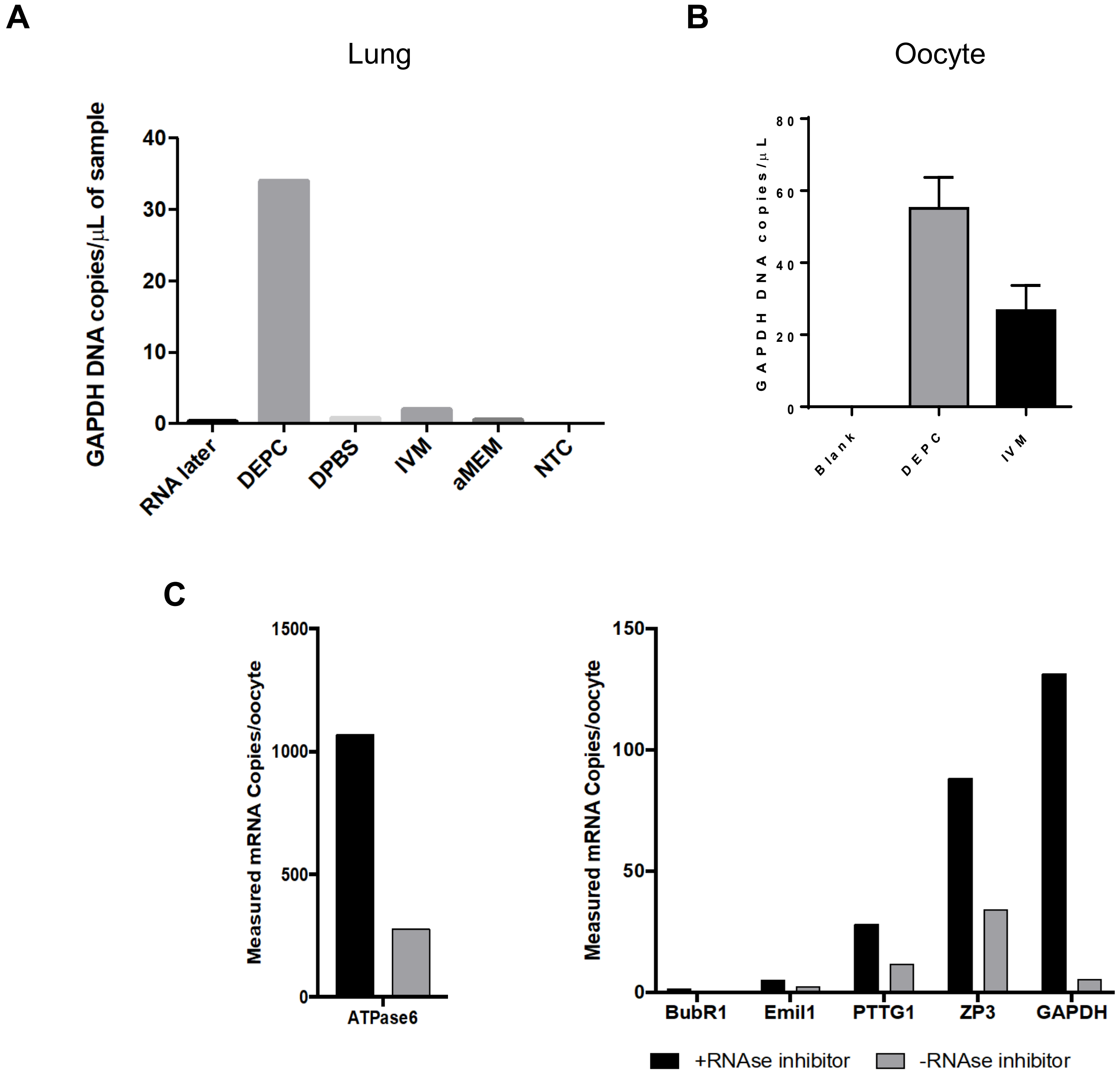

A
Lung
GAPDH DNA copies/µL of sample
40
30
20
10
0
RNA later
DEPC
DPBS
IVM
aMEM
NTC
B
Oocyte
GAPDH DNA copies/µL
80
60
40
20
0
Blank
DEPC
IVM
C
Measured mRNA Copies/oocyte
1500
1000
500
0
ATPase6
Measured mRNA Copies/oocyte
150
100
50
0
BubR1
Emil1
PTTG1
ZP3
GAPDH
+RNAse inhibitor
-RNAse inhibitor

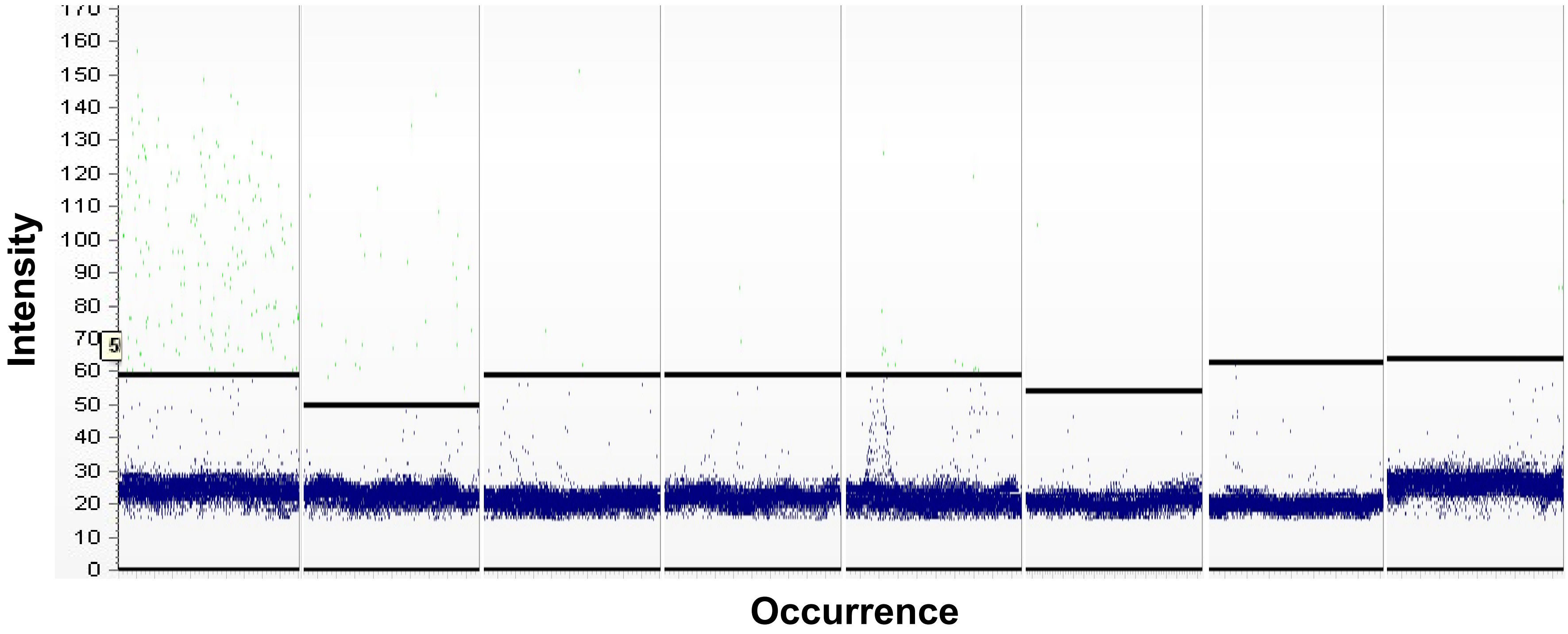

Intensity
170
160
150
140
130
120
110
100
90
80
70
60
50
40
30
20
10
0
5
Occurrence

**Table S1. PCR components**

**(i) mRNA analysis**

| Component | Final Concentration | Volume (µl) |
|---|---|---|
| 20X JN Solution | 1X | 0.75 |
| ATPase6/BuBR1/Emil1/PTTG1/ ZP3/Gapdh forward primers (10 µM) | 150nM | 0.225 |
| ATPase6/BuBR1/Emil1/PTTG1/ ZP3/Gapdh reverse primers (150 µM) | 150nM | 0.225 |
| 100X QuantiNova RT Mix | 1X | 0.15 |
| 2X QuantiNova SYBR Green RT-PCR Master Mix | 1X | 7.5 |
| Oocyte sample | | 6 |
| PCR Grade Water | | 0.15 |
| **Total** | | **15** |

**(ii) cDNA analysis**

| Component | Final Concentration | Volume (µl) | Volume (µl) |
|---|---|---|---|
| 20X JN Solution | 1X | 0.75 | 0.75 |
| Gapdh forward primers (10 µM) | 500nM | 0.75 | 0.75 |
| Gapdh reverse primers (10 µM) | 500nM | 0.75 | 0.75 |
| 2X SYBR Green | 1X | 7.5 | 7.5 |
| Mouse lung sample | | 3 | - |
| Oocyte sample | | - | 6 |
| PCR Grade Water | | 2.25 | - |
| **Total** | | **15** | **15.75** |

**(iii) Probe cDNA**

| Component | Final Concentration | Volume (µl) | Volume (µl) |
|---|---|---|---|
| 20X JN Solution | 1X | 0.75 | 0.75 |
| ATPase6/Gapdh forward primers (10 µM) | 250nM | 0.375 | 0.375 |
| ATPase6/Gapdh reverse primers (10 µM) | 250nM | 0.375 | 0.375 |
| ATPase6/Gapdh probe (50 µM) | 500nM | 0.15 | 0.15 |
| 2X QuantiNova Probe RT-PCR Master Mix | 1X | 7.5 | 7.5 |
| Mouse lung sample | | 3 | - |
| Oocyte sample | | - | 6 |
| PCR Grade Water | | 1.95 | - |
| **Total** | | **15** | **15.75** |

**(iv) RT**

| Reagents | Brand | Final Concentration | Volume |
|---|---|---|---|
| M-MLV RT 5x Reaction Buffer | Promega | 1x | 4 |
| dNTP Mix (5mM) | LifeTech | 0.5 mM | 2 |

| | | | |
|---|---|---|---|
| Random primers (0.5 µg/µL) | Promega | 1.0 µg/rxn | 1 |
| RNasin Ribonuclease Inhibitor (40 U/µL) | Promega | 20 U/rxn | 0.5 |
| M-MLV RT (200 U/µL) | Promega | 200 U/rxn | 1 |
| sample | | | 5 |
| $H_20$ | | - | 6.5 |
| **Total** | | | **20** |

**(v) miRNA RT**

| Component | Final Concentration | Volume (µl) |
|---|---|---|
| 5x RT Reaction Buffer | 1x | 2.86 |
| 10x RT Enzyme Mix | 1x | 1.43 |
| Sample | | 10 |
| **Total** | | **14.29** |

**(vi) miRNA cDNA dPCR**

| Component | Final Concentration | Volume (µl) |
|---|---|---|
| 2X SYBR Green | 1x | 7.5 |
| 20X JN Solution | 1x | 0.75 |
| Forward Primer | 150nM | 0.225 |
| Reverse Primer | 150nM | 0.225 |
| miRNA cDNA | | 6.3 |
| **Total** | | **15** |

**Table S2. Primer sequences**

**Messenger RNA Primer Sequences**

| Target | Forward Primer | Reverse Primer |
|---|---|---|
| GAPDH | GCAAGAGAGGCCCTATCCCAA | CTCCCTAGGCCCCTCCTGTTATT |
| ATPase6 | ACACCAAAAGGACGAACATGAA | TTCCTTGTGGAAGGAAGTGG |
| HPRT1 | GCTTGCTGGTGAAAAGGACCTCTCGAAG | CCCTGAAGTACTCATTATAGTCAAGGGCAT |
| Emil1 | GTGGAGGTGGCAAAGACATT | GGCAAAGGACCCACTTTACA |
| BubR1 | GCCAGGGAGAGAAGAAGG | CTCTCAGACTTGCAGTTATTTC |
| PTTG1 | CGAGTCGGCAAAGTGTTCAA | GCTCCCCAGGCAGGTC |
| ZP3 | CAGCCATGGCAACTGTAGTA | GCCTAACCCAAGAGCCAC |

**MicroRNA Primer Sequences**

| Target | Synthetic miRNA Sequence | Forward Primer | Reverse Primer |
|---|---|---|---|
| mmu-miR-485-5p | 5'- rArGrArGrGrCrUrGrGrCrCrGrUrGrArUrGrArArUrUrC -3' | GGCTGGCCGTGATGA | AGGTCCAGTTTTTTTTTTTTTTTGAA |
| mmu-miR-294-5p | 5'- rArCrUrCrArArArArUrGrGrArGrGrCrCrCrUrArUrCrU -3' | CGCAGACTCAAAATGGAG | AGGTCCAGTTTTTTTTTTTTTTTAGATAG |
| mmu-miR-183-5p | 5'- rUrArUrGrGrCrArCrUrGrGrUrArGrArArUrUrCrArCrU -3' | GCAGTATGGCACTGGTAGA | GGTCCAGTTTTTTTTTTTTTTTAGTGA |
| mmu-miR-196a-5p | 5'- rUrArGrGrUrArGrUrUrUrCrArUrGrUrUrGrUrUrGrGrG -3' | GCAGTAGGTAGTTTCATGTTGT | GTCCAGTTTTTTTTTTTTTTTCCCA |

**Table S3. Summary of (A) PCR chacteristics of GAPDH and (B) comparison of expected and observed E. coli GapA concentration as measured using dPCR.**

**(A)**

| Total cDNA (ng/µL) | Positive droplet number | Measured copies/µL of sample* | Ct | |
|---|---|---|---|---|
| 100 | Above upper limit of detection | | 20.68 | |
| 10 | 170 | 33 | 24.3945 | 425 |
| 1 | 30 | 5.53 | 27.1613 | 75 |
| 0.1 | 3 | 0.65 | 30.7435 | 7.5 |
| 0.01 | 4 | 0.78 | 36 | 10 |
| Control | 2 | 0.4 | - | |

*Measured copies/ul in sample = measured copies/ul in reaction mix × 15/6 (total reaction volume/DNA used)

**(B)**

| Expected concentration (copies/µL) | Mean of measured concentration (copies/µL) | Standard deviation | Relative standard deviation (%) | Difference(%) |
|---|---|---|---|---|
| 1800 | 1749.42 | 43.07 | 2.46 | -2.81 |
| 900 | 917.07 | 48.81 | 5.32 | 1.90 |
| 450 | 403.66 | 29.33 | 7.27 | -10.30 |
| 225 | 242.49 | 22.76 | 9.39 | 7.77 |
| 112.5 | 116.60 | 10.07 | 8.64 | 3.64 |

**Table S4. Extracellular microRNA by GV oocytes incubated with C24:1 Ceramide**

**Up-regulated**

| | miRNA | log2 Fold Change | Aveage Expression | P value | Adjusted p-value |
|---|---|---|---|---|---|
| 1 | mmu-miR-592-3p | 4.0781 | 2.5515 | 0.0114 | 0.0384 |
| 2 | mmu-miR-714 | 2.5484 | -4.0543 | 0.0215 | 0.0592 |
| 3 | mmu-miR-124-3p | 2.5389 | 2.0678 | 0.0439 | 0.0986 |
| 4 | mmu-miR-92b-5p | 2.3536 | 1.3018 | 0.0048 | 0.0243 |
| 5 | mmu-miR-1a-1-5p | 2.3130 | 3.7017 | 0.0029 | 0.0171 |
| 6 | mmu-miR-1931 | 2.2990 | 2.4299 | 0.0007 | 0.0069 |
| 7 | mmu-miR-500-3p | 2.1013 | -4.2258 | 0.0140 | 0.0445 |

**Down-regulated**

| 1 | miRNA | log2 Fold Change | Aveage Expression | P value | Adjusted p-value |
|---|---|---|---|---|---|
| 2 | mmu-miR-653-5p | -9.7507 | -1.4080 | 0.0000 | 0.0002 |
| 3 | mmu-miR-133a-3p | -7.6958 | -2.9457 | 0.0000 | 0.0002 |
| 4 | mmu-miR-485-5p | -7.4795 | -5.7043 | 0.0000 | 0.0003 |
| 5 | mmu-miR-196b-5p | -7.3789 | -0.7185 | 0.0000 | 0.0003 |
| 6 | mmu-miR-10b-3p | -7.1676 | -2.2734 | 0.0000 | 0.0004 |
| 7 | mmu-miR-489-3p | -6.9593 | -0.6417 | 0.0000 | 0.0005 |
| 8 | mmu-miR-431-5p | -6.7776 | -1.1712 | 0.0000 | 0.0005 |
| 9 | mmu-miR-145a-3p | -6.6963 | -0.0887 | 0.0000 | 0.0002 |
| 10 | mmu-miR-466g | -6.6650 | -0.8612 | 0.0000 | 0.0012 |
| 11 | mmu-miR-362-5p | -6.5690 | -5.0841 | 0.0000 | 0.0002 |
| 12 | mmu-miR-183-5p | -6.4754 | -3.0325 | 0.0000 | 0.0004 |
| 13 | mmu-miR-721 | -6.2614 | 0.3366 | 0.0000 | 0.0003 |
| 14 | mmu-miR-208b-3p | -6.2571 | 0.3388 | 0.0000 | 0.0002 |
| 15 | mmu-miR-15b-3p | -6.1058 | -0.1226 | 0.0000 | 0.0005 |
| 16 | mmu-miR-200c-5p | -5.9717 | -0.8423 | 0.0001 | 0.0014 |
| 17 | mmu-miR-196a-5p | -5.9563 | -3.6546 | 0.0000 | 0.0005 |
| 18 | mmu-miR-295-3p | -5.8322 | 0.5128 | 0.0000 | 0.0002 |
| 19 | mmu-miR-3062-5p | -5.7011 | -2.9661 | 0.0188 | 0.0534 |
| 20 | mmu-miR-466f-5p | -5.6860 | 0.0724 | 0.0001 | 0.0022 |
| 21 | mmu-miR-875-5p | -5.6847 | 0.6250 | 0.0000 | 0.0003 |
| 22 | mmu-miR-421-3p | -5.6820 | 0.3728 | 0.0000 | 0.0002 |
| 23 | mmu-miR-375-3p | -5.6339 | -1.0354 | 0.0000 | 0.0003 |
| 24 | mmu-miR-294-5p | -5.5172 | -4.2613 | 0.0011 | 0.0085 |
| 25 | mmu-miR-100-5p | -5.4368 | -4.0606 | 0.0000 | 0.0006 |
| 26 | mmu-miR-155-5p | -5.3388 | -0.1816 | 0.0001 | 0.0014 |
| 27 | mmu-miR-186-3p | -5.2768 | -3.6719 | 0.0001 | 0.0014 |
| 28 | mmu-miR-190a-5p | -5.2516 | 0.6453 | 0.0000 | 0.0003 |
| 29 | mmu-miR-181a-2-3p | -5.2468 | 0.8439 | 0.0000 | 0.0002 |
| 30 | mmu-miR-504-5p | -5.1518 | 0.2541 | 0.0001 | 0.0014 |
| 31 | mmu-miR-708-5p | -5.1161 | 0.4220 | 0.0000 | 0.0010 |
| 32 | mmu-miR-654-3p | -5.1024 | -1.6632 | 0.0025 | 0.0153 |
| 33 | mmu-miR-878-3p | -4.9439 | -0.8931 | 0.0021 | 0.0131 |
| 34 | mmu-miR-141-5p | -4.7257 | 1.1045 | 0.0000 | 0.0003 |
| 35 | mmu-miR-181d-5p | -4.6584 | -1.8145 | 0.0002 | 0.0024 |
| 36 | mmu-miR-291a-5p | -4.6101 | 0.2515 | 0.0007 | 0.0065 |
| 37 | mmu-miR-433-3p | -4.5814 | -2.1036 | 0.0000 | 0.0013 |
| 38 | mmu-miR-152-5p | -4.5536 | 0.2200 | 0.0002 | 0.0034 |
| 39 | mmu-miR-671-3p | -4.5272 | -0.0166 | 0.0001 | 0.0024 |
| 40 | mmu-miR-29a-3p | -4.3342 | -5.3104 | 0.0000 | 0.0006 |
| 41 | mmu-miR-804 | -4.3306 | 1.3020 | 0.0000 | 0.0004 |
| 42 | mmu-miR-201-5p | -4.1604 | 1.3871 | 0.0001 | 0.0014 |
| 43 | mmu-miR-770-3p | -4.1222 | 0.8259 | 0.0001 | 0.0014 |
| 44 | mmu-miR-135a-5p | -4.1174 | 0.7551 | 0.0001 | 0.0022 |
| 45 | mmu-miR-106b-3p | -4.0606 | 0.7242 | 0.0006 | 0.0060 |
| 46 | mmu-miR-181b-5p | -4.0509 | -1.3295 | 0.0000 | 0.0011 |

| | | | | | |
|---|---|---|---|---|---|
| 47 | mmu-miR-509-5p | -4.0068 | -2.2006 | 0.0009 | 0.0077 |
| 48 | mmu-miR-490-3p | -3.9783 | -1.5155 | 0.0009 | 0.0075 |
| 49 | mmu-miR-200b-5p | -3.8632 | 0.9133 | 0.0000 | 0.0008 |
| 50 | mmu-miR-380-5p | -3.8483 | -1.9836 | 0.0001 | 0.0018 |
| 51 | mmu-miR-302a-3p | -3.7795 | 0.0676 | 0.0001 | 0.0019 |
| 52 | mmu-miR-380-3p | -3.7476 | 0.8711 | 0.0001 | 0.0018 |
| 53 | mmu-miR-666-3p | -3.7186 | 1.2445 | 0.0002 | 0.0026 |
| 54 | mmu-miR-34b-3p | -3.6646 | -1.8305 | 0.0255 | 0.0662 |
| 55 | mmu-miR-135b-5p | -3.6118 | 1.6614 | 0.0001 | 0.0022 |
| 56 | mmu-miR-698-3p | -3.5605 | 1.3533 | 0.0006 | 0.0062 |
| 57 | mmu-miR-1969 | -3.5399 | -1.8829 | 0.0001 | 0.0014 |
| 58 | mmu-miR-421-5p | -3.5380 | -1.5886 | 0.0001 | 0.0022 |
| 59 | mmu-miR-667-3p | -3.5380 | 1.5916 | 0.0000 | 0.0013 |
| 60 | mmu-miR-101a-5p | -3.5213 | 1.7067 | 0.0001 | 0.0014 |
| 61 | mmu-miR-216b-5p | -3.5010 | 1.7168 | 0.0000 | 0.0010 |
| 62 | mmu-miR-9-5p | -3.4912 | 0.3875 | 0.0009 | 0.0073 |
| 63 | mmu-miR-326-3p | -3.4821 | 0.2388 | 0.0012 | 0.0087 |
| 64 | mmu-miR-872-5p | -3.4729 | 1.7309 | 0.0001 | 0.0021 |
| 65 | mmu-miR-148a-3p | -3.4519 | 1.7414 | 0.0003 | 0.0043 |
| 66 | mmu-miR-763 | -3.4281 | 0.6528 | 0.0010 | 0.0077 |
| 67 | mmu-miR-467c-5p | -3.4187 | 1.7580 | 0.0005 | 0.0058 |
| 68 | mmu-miR-682 | -3.3620 | 1.1304 | 0.0022 | 0.0135 |
| 69 | mmu-miR-223-5p | -3.3504 | -0.5866 | 0.0002 | 0.0029 |
| 70 | mmu-miR-151-3p | -3.3188 | 0.6780 | 0.0094 | 0.0334 |
| 71 | mmu-miR-532-5p | -3.3013 | -1.5094 | 0.0005 | 0.0055 |
| 72 | mmu-miR-1897-5p | -3.3003 | 1.8172 | 0.0001 | 0.0019 |
| 73 | mmu-miR-105 | -3.2779 | 1.6966 | 0.0001 | 0.0014 |
| 74 | mmu-miR-214-3p | -3.2687 | -1.0119 | 0.0040 | 0.0210 |
| 75 | mmu-miR-302a-5p | -3.2667 | 1.8340 | 0.0001 | 0.0014 |
| 76 | mmu-miR-881-5p | -3.2331 | 1.5377 | 0.0002 | 0.0024 |
| 77 | mmu-miR-678 | -3.2134 | -7.4756 | 0.0002 | 0.0024 |
| 78 | mmu-miR-142-3p | -3.1798 | 0.9376 | 0.0040 | 0.0210 |
| 79 | mmu-miR-542-3p | -3.1678 | -2.2242 | 0.0002 | 0.0034 |
| 80 | mmu-miR-99b-3p | -3.1598 | -6.2095 | 0.0001 | 0.0022 |
| 81 | mmu-miR-190b-5p | -3.1567 | 1.8890 | 0.0000 | 0.0011 |
| 82 | mmu-miR-423-3p | -3.1288 | -0.9352 | 0.0009 | 0.0073 |
| 83 | mmu-miR-701-5p | -3.1266 | 1.9040 | 0.0002 | 0.0027 |
| 84 | mmu-miR-1191 | -3.1060 | 0.4381 | 0.0022 | 0.0135 |
| 85 | mmu-miR-194-5p | -3.0839 | 1.6587 | 0.0013 | 0.0093 |
| 86 | mmu-miR-582-5p | -3.0654 | 1.9346 | 0.0000 | 0.0014 |
| 87 | mmu-miR-501-5p | -3.0581 | 1.9383 | 0.0001 | 0.0017 |
| 88 | mmu-miR-679-5p | -3.0457 | 0.6257 | 0.0007 | 0.0065 |
| 89 | mmu-miR-670-5p | -3.0242 | -3.1142 | 0.0001 | 0.0014 |
| 90 | mmu-miR-324-5p | -3.0238 | 0.7738 | 0.0157 | 0.0480 |
| 91 | mmu-miR-187-5p | -2.9969 | 0.0228 | 0.0004 | 0.0055 |
| 92 | mmu-miR-1896 | -2.9937 | 1.9705 | 0.0014 | 0.0100 |
| 93 | mmu-miR-3059-3p | -2.9854 | -5.7088 | 0.0015 | 0.0103 |
| 94 | mmu-miR-669k-5p | -2.9639 | 1.2978 | 0.0003 | 0.0044 |
| 95 | mmu-miR-743a-3p | -2.9612 | -0.7164 | 0.0020 | 0.0129 |
| 96 | mmu-miR-880-3p | -2.9180 | 1.8777 | 0.0007 | 0.0066 |
| 97 | mmu-miR-291b-3p | -2.9178 | 2.0084 | 0.0003 | 0.0042 |
| 98 | mmu-miR-302d-3p | -2.9050 | 1.8625 | 0.0036 | 0.0196 |
| 99 | mmu-miR-344d-3p | -2.8992 | -3.5519 | 0.0004 | 0.0052 |
| 100 | mmu-miR-434-3p | -2.8870 | -2.3002 | 0.0282 | 0.0713 |
| 101 | mmu-miR-429-3p | -2.8801 | -0.1990 | 0.0160 | 0.0480 |
| 102 | mmu-miR-293-3p | -2.8781 | 0.7369 | 0.0011 | 0.0086 |
| 103 | mmu-miR-1933-5p | -2.8626 | 1.4475 | 0.0017 | 0.0112 |
| 104 | mmu-miR-10a-3p | -2.8541 | -4.7341 | 0.0008 | 0.0069 |

| | | | | | |
|---|---|---|---|---|---|
| 105 | mmu-miR-344d-3-5p | -2.8515 | 1.8221 | 0.0006 | 0.0060 |
| 106 | mmu-miR-382-5p | -2.8270 | 0.4710 | 0.0076 | 0.0297 |
| 107 | mmu-miR-592-5p | -2.7874 | 1.6745 | 0.0039 | 0.0207 |
| 108 | mmu-miR-3474 | -2.7650 | -1.1058 | 0.0006 | 0.0065 |
| 109 | mmu-miR-709 | -2.7649 | 0.6778 | 0.0039 | 0.0209 |
| 110 | mmu-miR-297a-5p | -2.7353 | -2.2549 | 0.0019 | 0.0121 |
| 111 | mmu-miR-126-5p | -2.7077 | -0.2336 | 0.0011 | 0.0087 |
| 112 | mmu-miR-374b-3p | -2.7027 | 1.9031 | 0.0005 | 0.0055 |
| 113 | mmu-miR-383-5p | -2.6830 | -1.8031 | 0.0006 | 0.0065 |
| 114 | mmu-miR-361-3p | -2.6635 | -0.3922 | 0.0004 | 0.0055 |
| 115 | mmu-miR-186-5p | -2.6589 | -0.9254 | 0.0003 | 0.0041 |
| 116 | mmu-miR-30d-3p | -2.6567 | 2.1387 | 0.0001 | 0.0022 |
| 117 | mmu-miR-344c-3p | -2.6273 | -5.0183 | 0.0015 | 0.0105 |
| 118 | mmu-miR-875-3p | -2.6240 | 2.1553 | 0.0005 | 0.0056 |
| 119 | mmu-miR-28c | -2.6187 | 1.2233 | 0.0007 | 0.0065 |
| 120 | mmu-miR-182-5p | -2.6143 | 2.0109 | 0.0002 | 0.0026 |
| 121 | mmu-miR-3063-3p | -2.5965 | 0.4089 | 0.0098 | 0.0345 |
| 122 | mmu-miR-451a | -2.5820 | -0.0466 | 0.0224 | 0.0608 |
| 123 | mmu-miR-28a-3p | -2.5722 | -0.6649 | 0.0009 | 0.0073 |
| 124 | mmu-miR-206-3p | -2.5672 | -0.5544 | 0.0002 | 0.0024 |
| 125 | mmu-miR-683 | -2.5601 | 2.1873 | 0.0008 | 0.0073 |
| 126 | mmu-miR-107-5p | -2.5490 | 1.6851 | 0.0011 | 0.0085 |
| 127 | mmu-miR-3112-3p | -2.5473 | 0.3457 | 0.0008 | 0.0069 |
| 128 | mmu-miR-297b-5p | -2.5440 | 0.7511 | 0.0068 | 0.0297 |
| 129 | mmu-miR-541-3p | -2.5412 | -0.3643 | 0.0005 | 0.0055 |
| 130 | mmu-miR-221-3p | -2.5368 | 0.1645 | 0.0159 | 0.0480 |
| 131 | mmu-miR-1960 | -2.5361 | -1.4948 | 0.0009 | 0.0075 |
| 132 | mmu-miR-30c-2-3p | -2.5351 | 0.9688 | 0.0300 | 0.0737 |
| 133 | mmu-miR-881-3p | -2.5239 | -11.0207 | 0.0004 | 0.0053 |
| 134 | mmu-miR-125b-1-3p | -2.5104 | 1.7198 | 0.0004 | 0.0051 |
| 135 | mmu-miR-761 | -2.5053 | 1.8880 | 0.0005 | 0.0055 |
| 136 | mmu-miR-3097-5p | -2.5032 | 0.3882 | 0.0103 | 0.0356 |
| 137 | mmu-miR-297c-5p | -2.4900 | 1.8174 | 0.0012 | 0.0087 |
| 138 | mmu-miR-181d-3p | -2.4893 | -7.0601 | 0.0005 | 0.0055 |
| 139 | mmu-miR-361-5p | -2.4825 | 1.0587 | 0.0032 | 0.0182 |
| 140 | mmu-miR-665-5p | -2.4622 | 1.2951 | 0.0350 | 0.0835 |
| 141 | mmu-miR-497-5p | -2.4562 | 2.2392 | 0.0081 | 0.0302 |
| 142 | mmu-miR-101b-5p | -2.4402 | 1.8297 | 0.0004 | 0.0055 |
| 143 | mmu-miR-3471 | -2.4333 | -4.7932 | 0.0192 | 0.0543 |
| 144 | mmu-miR-1945 | -2.4188 | -1.6547 | 0.0005 | 0.0058 |
| 145 | mmu-miR-196a-1-3p | -2.4175 | 1.7942 | 0.0036 | 0.0195 |
| 146 | mmu-miR-1902 | -2.4104 | 1.6310 | 0.0051 | 0.0252 |
| 147 | mmu-miR-574-3p | -2.4067 | 1.8733 | 0.0029 | 0.0171 |
| 148 | mmu-miR-302d-5p | -2.3836 | -1.8993 | 0.0014 | 0.0100 |
| 149 | mmu-miR-1947-5p | -2.3828 | 2.2759 | 0.0006 | 0.0065 |
| 150 | mmu-miR-331-3p | -2.3682 | -0.4870 | 0.0112 | 0.0378 |
| 151 | mmu-miR-3071-5p | -2.3656 | 0.0518 | 0.0030 | 0.0172 |
| 152 | mmu-miR-200a-5p | -2.3385 | -0.6457 | 0.0053 | 0.0261 |
| 153 | mmu-miR-10a-5p | -2.3308 | -0.0406 | 0.0009 | 0.0075 |
| 154 | mmu-miR-325-5p | -2.3144 | -0.1200 | 0.0013 | 0.0093 |
| 155 | mmu-miR-688 | -2.3079 | 2.3134 | 0.0017 | 0.0112 |
| 156 | mmu-miR-29c-5p | -2.3048 | -3.1331 | 0.0010 | 0.0077 |
| 157 | mmu-miR-207 | -2.3035 | -0.4319 | 0.0281 | 0.0713 |
| 158 | mmu-miR-146b-3p | -2.2946 | 2.3200 | 0.0034 | 0.0191 |
| 159 | mmu-miR-130b-3p | -2.2914 | -1.3304 | 0.0058 | 0.0274 |
| 160 | mmu-miR-544-5p | -2.2829 | 1.4551 | 0.0141 | 0.0445 |
| 161 | mmu-miR-3087-3p | -2.2555 | -0.1211 | 0.0009 | 0.0073 |
| 162 | mmu-miR-669c-5p | -2.2458 | -0.9882 | 0.0013 | 0.0094 |

| | | | | | |
|---|---|---|---|---|---|
| 163 | mmu-miR-93-3p | -2.2447 | -2.4184 | 0.0057 | 0.0274 |
| 164 | mmu-miR-26a-1-3p | -2.2323 | 0.1056 | 0.0015 | 0.0105 |
| 165 | mmu-miR-496a-5p | -2.2270 | -2.7279 | 0.0007 | 0.0065 |
| 166 | mmu-miR-3072-3p | -2.2246 | 2.3550 | 0.0007 | 0.0066 |
| 167 | mmu-miR-222-5p | -2.2146 | 0.0432 | 0.0050 | 0.0250 |
| 168 | mmu-miR-329-5p | -2.2122 | -1.6674 | 0.0054 | 0.0264 |
| 169 | mmu-miR-3095-5p | -2.2087 | 2.0012 | 0.0187 | 0.0534 |
| 170 | mmu-miR-1928 | -2.2077 | 1.9885 | 0.0008 | 0.0069 |
| 171 | mmu-miR-18b-5p | -2.1982 | 0.3998 | 0.0161 | 0.0480 |
| 172 | mmu-miR-411-5p | -2.1845 | 2.3751 | 0.0174 | 0.0509 |
| 173 | mmu-miR-702-3p | -2.1786 | 0.4359 | 0.0148 | 0.0460 |
| 174 | mmu-miR-693-3p | -2.1772 | 1.1475 | 0.0088 | 0.0319 |
| 175 | mmu-miR-96-5p | -2.1683 | 1.5826 | 0.0098 | 0.0346 |
| 176 | mmu-miR-224-5p | -2.1634 | 2.3856 | 0.0079 | 0.0301 |
| 177 | mmu-miR-485-3p | -2.1525 | -0.8147 | 0.0160 | 0.0480 |
| 178 | mmu-miR-695 | -2.1493 | -10.6637 | 0.0027 | 0.0160 |
| 179 | mmu-miR-92b-3p | -2.1481 | -3.3075 | 0.0005 | 0.0057 |
| 180 | mmu-miR-539-3p | -2.1477 | 1.4919 | 0.0243 | 0.0640 |
| 181 | mmu-miR-340-5p | -2.1469 | 0.8716 | 0.0174 | 0.0509 |
| 182 | mmu-miR-1251-3p | -2.1376 | 2.2139 | 0.0013 | 0.0096 |
| 183 | mmu-miR-463-3p | -2.1363 | -3.8420 | 0.0035 | 0.0192 |
| 184 | mmu-miR-376a-3p | -2.1286 | 2.4030 | 0.0080 | 0.0301 |
| 185 | mmu-miR-1948-5p | -2.1276 | 1.5825 | 0.0081 | 0.0302 |
| 186 | mmu-miR-381-3p | -2.1021 | 2.2136 | 0.0012 | 0.0088 |
| 187 | mmu-miR-3061-3p | -2.0957 | 2.1289 | 0.0006 | 0.0061 |
| 188 | mmu-miR-1912-5p | -2.0850 | 0.2829 | 0.0487 | 0.1067 |
| 189 | mmu-miR-130b-5p | -2.0594 | 2.4376 | 0.0011 | 0.0085 |
| 190 | mmu-let-7d-3p | -2.0537 | 0.2845 | 0.0055 | 0.0266 |
| 191 | mmu-miR-344g-5p | -2.0295 | -0.3446 | 0.0102 | 0.0356 |
| 192 | mmu-miR-182-3p | -2.0256 | 1.4757 | 0.0110 | 0.0373 |
| 193 | mmu-miR-1970 | -2.0239 | 2.4554 | 0.0017 | 0.0110 |
| 194 | mmu-miR-3105-3p | -2.0234 | 2.1708 | 0.0167 | 0.0495 |
| 195 | mmu-miR-764-5p | -2.0153 | -2.1555 | 0.0010 | 0.0083 |
| 196 | mmu-miR-1264-3p | -2.0119 | 1.0154 | 0.0385 | 0.0894 |
| 197 | mmu-miR-1930-3p | -2.0041 | -0.4641 | 0.0126 | 0.0412 |
| 198 | mmu-miR-711 | -2.0037 | 2.2661 | 0.0064 | 0.0293 |
| 199 | mmu-miR-676-5p | -2.0015 | 2.3786 | 0.0007 | 0.0065 |
